# Non-operable glioblastoma: proposition of patient-specific forecasting by image-informed poromechanical model

**DOI:** 10.1101/2023.01.10.523475

**Authors:** Stéphane Urcun, Davide Baroli, Pierre-Yves Rohan, Wafa Skalli, Vincent Lubrano, Stéphane P.A. Bordas, Giuseppe Sciumè

## Abstract

We propose a novel image-informed glioblastoma mathematical model within a reactive multiphase poromechanical framework. Poromechanics offers to model in a coupled manner the interplay between tissue deformation and pressure-driven fluid flows, these phenomena existing simultaneously in cancer disease. The model also relies on two mechano-biological hypotheses responsible for the heterogeneity of the GBM: hypoxia signaling cascade and interaction between extra-cellular matrix and tumor cells. The model belongs to the category of patient-specific image-informed models as it is initialized, calibrated and evaluated by the means of patient imaging data. The model is calibrated with patient data after 6 cycles of concomitant radiotherapy chemotherapy and shows good agreement with treatment response 3 months after chemotherapy maintenance. Sensitivity of the solution to parameters and to boundary conditions is provided. As this work is only a first step of the inclusion of poromechanical framework in image-informed glioblastoma mathematical models, leads of improvement are provided in the conclusion.

## 1. Introduction

In global cancer statistics, primary brain tumors hold the 21^st^ rank of incidence and reach the 14^th^ rank of mortality [1]. Glioma represent the majority of malignant primary brain tumor. The group of diffuse glioma - ‘diffuse’ being opposed to ‘circumscribed’ - has the worst prognosis. The diagnosis of diffuse glioma was first based on histological features as infiltrative glioma cells along pre-existing tissue elements, historically known as secondary Scherer’s structures. In 2016, the previous classification the World Health Organisation (WHO) of diffuse glioma [2] was based on specific and cumulative histological features: nuclear atypia for grade 2, mitotic activity for grade 3 denoted anaplasic, necrosis and/or microvascular proliferation for grade 4, denoted glioblastoma multiforme (GBM). This classification also included molecular biomarkers, and specifically the isocitrate dehydrogenase (IDH) status. The IDH status was significant enough to lead to the new 2021 WHO classification [3], where an IDH wild-type status is directly classified a grade 4 GBM. Consequently, the GBM is now defined by this IDH wild-type status, and the IDH mutated status is termed as Astrocytoma, from grade 1 to 3. Grade 4 has the poorest prognosis with a median survival around 15 months, and a 5-year survival rate at 5.8%, constant since the year 2000 [4]. In this article, we are specifically interested in the IDH wild-type status. In the clinical literature, between 16% and 40% of GBM are considered non operable [5, 6], because of a functional critical location which impedes the resection, or because of patient co-morbidity. Non-operable cases allow longitudinal data of glioblastoma evolution, on a patient specific basis. Hence, they are of critical interest for modeling and forecasting processes.

GBM have received a large attention from the modeling community. An review of glioblastoma modeling was made by Falco *et al*. in [7] in 2021 and Mang *et al*. in [8] in 2020, the latter specifically focusing on image-informed glioblastoma modeling. New hypotheses may emerge from *in silico* studies and treatment personalization may be facilitated by the exploration *in silico* of the parameter space of the patient. This highly lethal disease and the absence of improvement of its survival rate made these two challenges particularly urgent. The authors of [7] reviewed 295 articles published between 2001 and 2020, and defined three categories, continuous, discrete and hybrid. The continuous models considered the disease as a collection of tissue and the targets of this type of model is the invasion pattern and the treatment response at the macroscale (for instance, see [9, 10]). The discrete models are tailored for the description of intra-cellular phenomena and interaction at the cellular level. They target genetic and immunological properties. Hybrid models try to retrieve the best of both approaches, by informing the models with multi-scale data, such as histological staining, genetic markers and clinical imaging (for discrete and hybrid categories, see [11, 12]). Some of these models, whatever their category, may be initialized and calibrated by clinical imaging data. By this means, they aim to patient-specific results. This supra-category is termed as image-informed model. This modeling framework was first developed in 2002, and applied to low and high grade gliomas, by Swanson *et al*. in [13], and after in [14, 15, 16]. Since 2013, with the progress of imaging methods, this framework has been further developed by Yankeelov *et al*. (see [17]), and also applied with clinically-relevant results in various locations such as breast cancer [18] or prostate cancer [19]. Image-informed glioblastoma modeling have been extensively used in the last decades, and have led to personalized modeling in tumor forecasting and treatment response [20, 21, 22], and to the inclusion of tissue anisotropy [23], among others hypotheses.

We propose in this article a novel image-informed glioblastoma model within a continuous multiphase poromechanical framework. Poromechanics offers to model the coupling between tissue deformation and pressure-driven fluid flows, these phenomena existing simultaneously in cancer disease. Poromechanics is already applied in cancer modeling, *in vitro* [24, 25] and in animal model [26]. However, except for a proposition of patient-specific image-informed modeling in [27] with only qualitative results, to our knowledge, there is no example of this framework applied to glioblastoma modeling in a clinically-relevant and patient-specific basis. Additionally to the description of brain tissue as a porous medium, our model relies on two mechano-biological hypotheses responsible for the heterogeneity of the GBM: hypoxia signaling cascade [28] and interaction between extra-cellular matrix and tumor cells [29]. A subset of the parameters of the model is initialized with the first time point of the patient imaging data, performed during pre-operative examination. The patient being non-operable, the simulation outputs are calibrated against patient’s imaging after only 6 cycles of concomitant radiotherapy-temozolomide chemotherapy (RT-TMZ), performed 63 days after the initial time. Through patient’s segmentations, the quantity evaluated are the overlapping between the clinical and the numerical tumors. After this calibration, the results are validated against a patient follow-up imaging 165 days after the initial time.

In the article, we briefly present the GBM and its management, followed by the presentation of the mathematical model, the patient dataset, the calibration process and a preliminary evaluation of the simulation. The results section gives the solution sensitivity on parameters variation and error of the model measured against patient imaging. Mathematical verification, such as solution sensitivity on boundary conditions are provided. As this work is only a first step of the inclusion of poromechanics in image-informed GBM modeling, we discuss the improvements and further propositions for this inspiring modeling framework.

### Description of the GBM according to the WHO 2021 classification

Glioma may originate from three sources [30]:

- neural stem cells, embryonic cells located in ventricular and subventricular zones of the brain, which give rise to both neurons and glial cells;
- oligodendrocyte precursor cells, a subset of glial cells precursor specific to oligodendrocytes;
- astrocyte, for which a specific precursor is not yet identified.

Therefore, the origin of the cellular population, and of the mutations in this population, that give rise to glioma, remains open for debate [30]. However, already developed GBM always have an astrocytic profile. This profile is characterized by a high heterogeneity both genetic and phenotypic, which creates difficulties both in origin determination and therapeutic design. Among diffuse glioma, GBM is by far the most common (90%). They are the majority of glioma and almost predominant among primary malignant brain tumors. The median age at diagnosis is 65 years and the male incidence is 50% higher than female. Except for radiation and rare genetic syndromes, there is no validated risk factor. Since 2005, its standard of care is, if possible, surgical resection followed by six 1-week cycles of concomitant radiotherapy and temozolomide chemotherapy [31], denoted RT-TMZ treatment. The TMZ is used as a radio sensitizer, and after the 6 cycles, TMZ only is used as maintenance from six to twelve months. Despite improvement of the median survival, now *>* 15 months, glioblastoma still have a poor prognosis, with a 5-year survival rate at 5.8%, constant since 2000 [4].

The 2016 WHO classification included molecular biomarkers, which previously defined GBM subtypes. The first biomarker was the status of isocytrate dehydrogenase (IDH). Non-mutated, the subtype is termed wild-type, other subtypes are mutated IDH-1 or IDH-2. IDH 1, 2 or 3 are enzymes involved in cell metabolism. If the 2016 WHO classification admitted two GBM subtypes, IDH wild-type and IDH mutated, the new 2021 WHO classification only considers one type of GBM, the IDH wild-type. The main reason is IDH mutated are lower grade astrocytomas that evolved into a higher grade, where IDH wild-type are of high grade since the diagnosis. Therefore, GBM IDH wild-type are now simply termed GBM, and WHO 2016 GBM IDH mutated are now termed astrocytomas grade 4. Mutation of IDH 1 or 2, by disrupting DNA demethylation, lead to the accumulation of an inhibitor of glioma stem cell differentiation [32]. However, IDH mutants represented 10% of all glioma grade 4 and have a better prognosis, as they are less resistant to chemotherapy and provoke better immune response [33]. A second marker is the status of the O^6^-methylguanine–DNA methyltransferase (MGMT), mythelated or non-methylated. The MGMT gene encodes a DNA-repair protein, therefore a high MGMT activity in cancer creates a resistant phenotype both on chemo- and radiotherapy. MGMT activity can be silenced by methylation and it decreases the DNA-repair activity [34]. The methylation of MGMT represents around 25% of GBM cases. This marker will influence the patient response to the RT-TMZ treatment, as a methylated MGMT profile is considered be more sensitive to RT effect [35]. The IDH status and the MGMT methylation status are not correlated, both types of markers can co-exist.

A necrotic core and/or an abnormal micro-vasculature are always present in GBM. These characteristics indicate that hypoxia management is a key feature of GBM. Barnes *et al*. show in [29] that hypoxia applied on GBM cells provokes structural changes on the surrounding ECM. The brain ECM has a specific composition. Conversely to the usual rich fibrillar component such as collagen, brain ECM is almost entirely composed of glycosaminoglycans, a non-fibrous component which plays the mechanical role of shock-absorber. GBM cells subjected to a hypoxic environment modify the structure of glycosaminoglycans. Hypoxia signaling is made through hypoxia-inducible factor-1*α* (HIF1*α*), which provokes the production by the GBM cells of the glycoprotein tenascin-C. The tenascin-C modifies the surronding glycosaminoglycans, leading to a cross-linked, stiffer ECM. In this article, we hypothesize that this ECM stiffening coupled with proliferative GBM cells will ultimately lead to an environment with a higher mechanical stress. Conversely, glioma cells with a IDH mutated status have a reduced capacity to produce both HIF1*α* and tenascin-C [36]. Therefore, this high stiffness of the tumorous tissue is characteristic of GBM.

#### 2. Reactive poromechanical modeling of GBM IDH wild-type

The model presented in this section belongs to the category termed as image-informed reactive multiphase poromechanics. Let us describe each part of this category:

- Poromechanics: the physical system is considered as a composite continuum composed of a permeable and deformable solid scaffold in which and through which fluid flows.
- multiphase: solid and fluid compartments are composite. The solid fraction, which could be compared to the medical definition of the stroma, is made of different and distinct materials (epithelial tissue, ECM - itself composite -, wall vessels, to name a few). Likewise, the fluid fraction is composed of different phases (interstitial fluid, immune cells, tumor cells). It should be noted that the blood is not modeled as a circulating fluid in this model.
- Reactive: the modeling of living tissue implies the biological interactions of many diffusive chemical agents (oxygen, cytokines), which can belong to any phase of the system. Their own dynamics are strongly coupled with the poromechanical system. The model also includes non-diffusive reactions such as mechanically-induced phenotype switch and hypoxic-induced necrosis.
- Image-informed: in order to simulate patient-specific cases, the initial conditions and the boundary conditions of the problem are provided by the patient MRI measurements. A subset of the model’s parameters is fixed by these measurements, another subset is calibrated with them.

### General framework

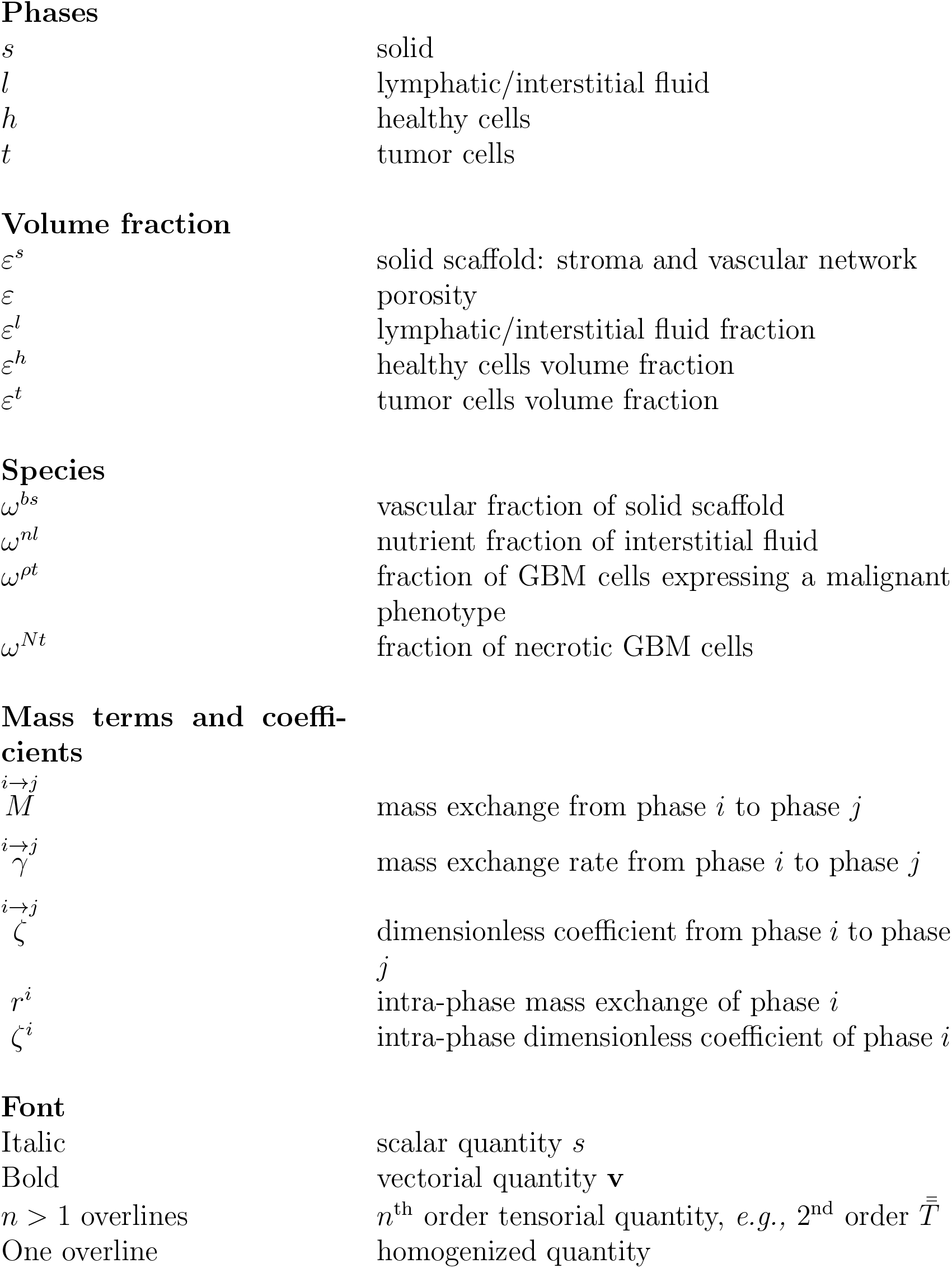

Phases
|  |  |
| --- | --- |
| $s$ | solid |
| $l$ | lymphatic/interstitial fluid |
| $h$ | healthy cells |
| $t$ | tumor cells |
| Volume fraction |  |
| $\varepsilon^s$ | solid scaffold: stroma and vascular network |
| $\varepsilon$ | porosity |
| $\varepsilon^l$ | lymphatic/interstitial fluid fraction |
| $\varepsilon^h$ | healthy cells volume fraction |
| $\varepsilon^t$ | tumor cells volume fraction |
| Species |  |
| $\omega^{bs}$ | vascular fraction of solid scaffold |
| $\omega^{nl}$ | nutrient fraction of interstitial fluid |
| $\omega^{\rho t}$ | fraction of GBM cells expressing a malignant phenotype |
| $\omega^{Nt}$ | fraction of necrotic GBM cells |
| Mass terms and coefficients |  |
| $M^{i \rightarrow j}$ | mass exchange from phase $i$ to phase $j$ |
| $\gamma^{i \rightarrow j}$ | mass exchange rate from phase $i$ to phase $j$ |
| $\zeta^{i \rightarrow j}$ | dimensionless coefficient from phase $i$ to phase $j$ |
| $r^i$ | intra-phase mass exchange of phase $i$ |
| $\zeta^i$ | intra-phase dimensionless coefficient of phase $i$ |
| Font |  |
| Italic | scalar quantity $s$ |
| Bold | vectorial quantity $\mathbf{v}$ |
| $n > 1$ overlines | $n^{\text{th}}$ order tensorial quantity, *e.g.*, 2<sup>nd</sup> order $\bar{\bar{T}}$ |
| One overline | homogenized quantity |

Let *ε*^*s*^, the volume fraction occupied by the solid scaffold and *ε*, the volume fraction occupied by the fluid phases.

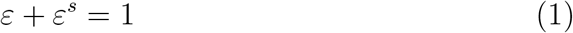

The vascular network *ω*^*bs*^ is considered as a fraction of the solid scaffold, its volume fraction is denoted *ε*^*s*^*ω*^*bs*^.

Considering the fluid phases (*t*, tumor, *h*, healthy and *l*, fluid) and defining their own saturation degree as *S*^*β*^ = *ε*^*β*^/*ε* (with *β* = *t, h, l* the index associated to extra-vascular fluids), we obtain:

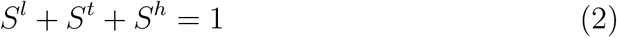

Their respective volume fraction are defined by *ε*^*β*^ = *εS*^*β*^.

To facilitate the understanding of the terms of the governing equations, we report the general form the mass conservation equations for a phase and a species provided by the thermodynamically constrained averaging theory (TCAT) [37] framework. The spatial form of the mass balance equation for an arbitrary phase *α* reads

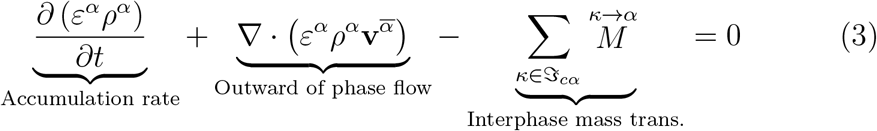

where *ρ*^*α*^ is the density, 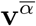 is the local velocity vector, 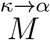 are the mass exchange terms accounting for transport of mass at the *κα* interface from phase *κ* to phase *α*, and 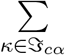 is the summation over all the phases sharing interfaces with the phase *α*.

An arbitrary species *i* dispersed within the phase *α* has to satisfy mass conservation too. The following spatial equation is derived following TCAT

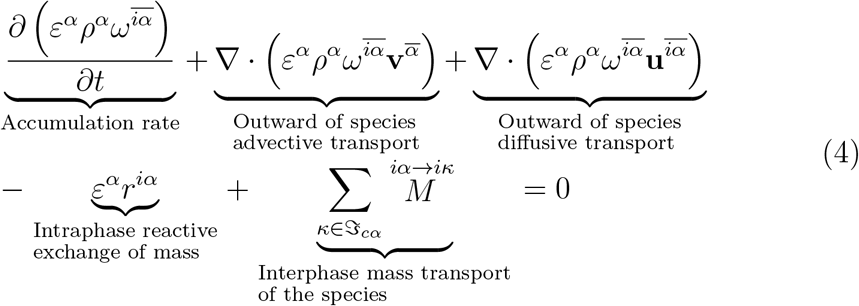

where 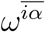 identifies the mass fraction of the species *i* dispersed with the phase *α, ε*^*α*^*r*^*iα*^ is a reaction term that allows to take into account the reactions between the species *i* and the other chemical species dispersed in the phase *α*, and 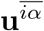 is the diffusive velocity of the species *i*.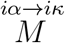 are mass exchange terms accounting for mass transport of species *i* at the *κα* interface from phase *α* to phase *κ*.

### Governing equations

The solid scaffold being deformable, we use the chain rule to define the material derivative:

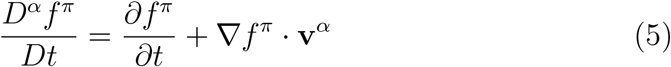

And apply it to Eq.3 and 4

We define the mass conservation of phases by using Eq.5 to express derivatives with respect to the solid phase *ε*^*s*^. Introducing porosity *ε* and the saturation degrees of its phases *t, h*, and *l*, the mass balance equations of *s, t, h* and *l* phases read respectively:

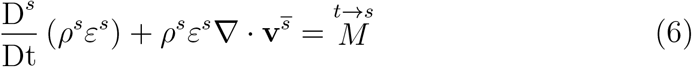

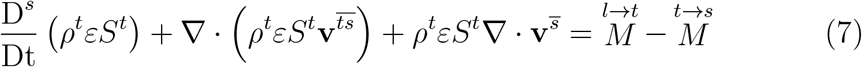

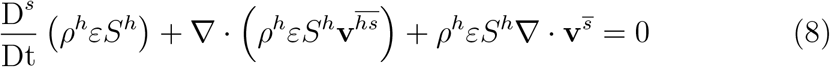

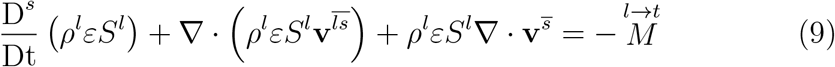

This system can be summarized as follows:

- tumorous phase takes its mass from interstitial fluid phase 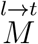;
- tumorous phase produce solid (fibrous) components 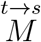;
- healthy cellular phase is considered at the equilibrium.

### Mass conservation equations of species

The only diffusive species considered is the oxygen, dissolved in the interstitial fluid phase *l*, its mass fraction denoted 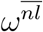. It motion is governed by advection-diffusion equation. The species is produced by micro-capillaries of the solid fraction phase *ω*^*bs*^, and absorbed by *t* and *h*, tumor and healthy cells, its mass balance reads

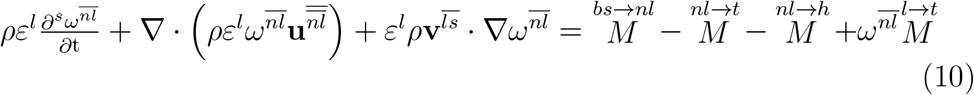

The necrotic fraction of tumor cells 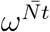 is a non-diffusive species. We obtain:

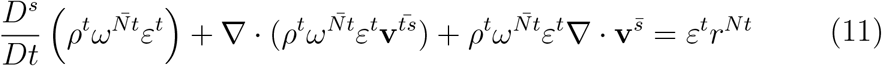

with the constitutive equation of the necrotic growth rate *r*^*Nt*^ (see Eq.40).

### Momentum equations

The porous system is modelled as a continuous medium, under linear momentum conservation:

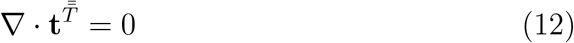

Where 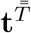 is the total Cauchy stress tensor. We assume here that all phases are incompressible. However, the overall multiphase system is not incompressible because, as an open system, the presence of porosity can evolve according to the scaffold deformation. As all phases are incompressible, their densities *ρ*^*α*^ (with *α* = *s, t, l*) are constant and the Biot’s coefficient *β* = 1. With these premises, the total Cauchy stress tensor appearing in Eq.12 is related to the Biot’s effective stress as follows

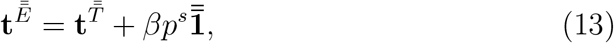

where *p*^*s*^ = *S*^*l*^*p*^*l*^ + *S*^*t*^*p*^*h*^ + *S*^*t*^*p*^*h*^ is denoted the solid pressure, describing the interaction between the fluids and the solid scaffold. From a clinical point of view, *p*^*s*^ corresponds to the intracranial pressure, with each component of the multiphase system contributing to the exerted pressure.

### Internal variables

#### ECM stiffening

One internal variable, the Young’s Modulus of the ECM *E*^*ECM*^, is updated every 250 minutes, *i*.*e*. every 10 iterations. This corresponds to a physical quantity that has a slower evolution than the primary unknowns (the displacement field, the pressures of the fluids and the level of oxygen). In the following equations, *T* = 250 min. The stiffening of the ECM is modelled as follows:

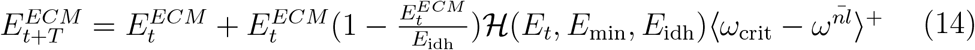

where *E*_min_ fixed at the lower bound of the stiffness measured in the cortex tissue *E*_min_ = 1.2kPa and *E*_idh_, the stiffness of cross-linked ECM, is to be calibrated. The regularized step function ℋ is used in several constitutive equations and given in Eq.34.

#### Malignant fraction and RT-TMZ treatment

Two other internal variables, the fraction of GBM cells expressing a malignant phenotype *ω*^*ρt*^ and the administration of the RT-TMZ treatment are updated on a daily basis. *ω*^*ρt*^ is updated every 4.5 days, which corresponds to 260 iterations. We note a lack of quantitative information about phenotype switch in the experimental literature. However, we found that at the cell scale, phenotype switch can be measured in minutes or in hours [38]. The only example we found at the macroscale is about lung cancer cells, where the effects of a phenotype switch is observable after a minimum of 72 hours [39]. In the absence of further information on GBM cells, we keep our range *T* = 4.5 days. If the tumorous region undergoes a high osmotic pressure,*i*.*e*. greater than the threshold *p*_idh_, and a chronic hypoxia, during the period *T*, a fraction of IDH wild-type cells *ω*^*ρt*^ changes their phenotype. This fraction is updated as follows:

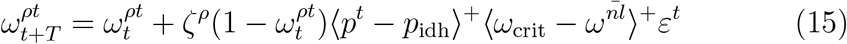

with *ζ*^*ρ*^ the phenotype switch rate, *ε*^*t*^ the volume fraction of GBM cells, and ⟨*α* − *β*⟩^+^ = 0 if *α < β* and 1 else.

The RT-TMZ treatment is administrated by following the standard of care defined in 2005 in [31]: before 1 month after diagnosis, the non-operable patient started 6 weekly cycles: 5 daily doses of 2 Gray radiotherapy (RT) concomitant with a daily dose of Temozolomide (TMZ). The second part of the standard treatment consists in 24 weeks of daily TMZ. The patient of this study has an non-methylated MGMT profile, which is more resistant to the RT-TMZ treatment [35]. The treatment is modeled by both long and short term effects. The short term effect provokes the necrosis of the tissues, preferentially the tumorous tissue. In this article, we only model the necrosis of GBM cells, with two dependencies. First, the TMZ being transmitted through the vascular network, its effect increases according to the local vascular fraction. Second, the denser regions of the tumor are known to have a more resistant profile [40]. These two dependencies are modeled by the following equations:

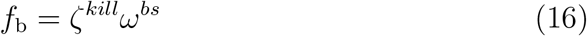

*f*_b_ represents the vascular dependency of RT-TMZ. *ζ*^*kill*^ is the optimal killing rate of cells by RT, *ω*^*bs*^ the vascular fraction of the solid scaffold.

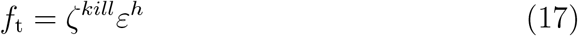

*f*_t_ represents the TC density dependency of RT-TMZ. *ε*^*h*^ is the volume fraction of healthy cells.

RT-TMZ short effect on the necrotic fraction 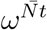 is modeled by the following equation, the period *T* = 1 day, 5 days per week:

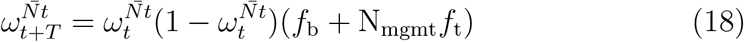

with N_mgmt_, the negative status of the methylation of MGMT.

The long effect, representative of the sole TMZ activity (6 RT-TMZ cycles plus 24 weeks of TMZ maintenance), is modeled on TC activity as follows:

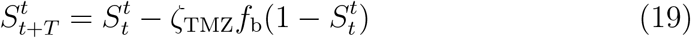

where *ζ*_TMZ_ quantify the patient’s response to TMZ.

#### Constitutive relationships

##### Stress-strain relationship

For the solid scaffold deformation, the chosen closure relationship for the effective stress 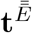 is linear elastic:

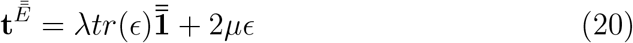

with 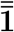 the identity tensor, 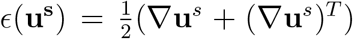 the linearised strain tensor, and the Lamé constant 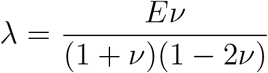 and 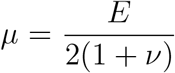. The two parameters *E* (except the stiffened *E*^ECM^) and *ν* were calibrated by *ex vivo* mechanical testing in [41].

#### Generalized Darcy’s law

The interaction between fluid phases and the solid scaffold are modeled by a generalized Darcy’s flow, deduced from the linear momentum conservation of fluid phases. The details of this constitutive relationship are provided in [42] and [25].

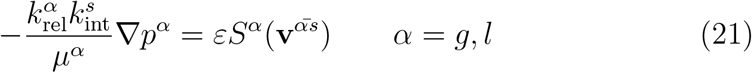

where 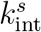 is the intrinsic permeability of the solid scaffold, *μ*^*α*^, 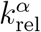 and *p*^*α*^ are respectively the dynamic viscosity, relative permeability and the pressure of each fluid phase *α* = *l, h, t*. The three fluid phases have their own relative permeabilities:

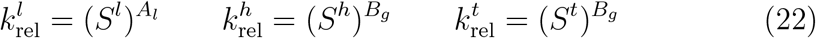

with *A*_*l*_ and *B*_*g*_, the exponents governing the evolution of the relative permeabilities. Both were calibrated by *ex vivo* mechanical testing in [41]. We choose to apply *B*_*g*_ to both glial and glioma cells.

#### Pressure-saturation relationships

The porosity is saturated by three immiscible fluid phases. Each phase has its own pressure. Three capillary pressures *p*^*ij*^, *i*.*e*. pressure difference between fluid *i* and fluid *j*, can be defined

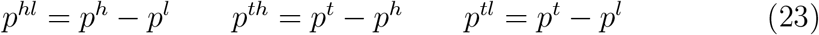

As in [43], we assume here that IF is the wetting fluid, HC is the intermediate-wetting fluid and TC the non-wetting fluid. Only two between the previously defined capillary pressures are independent since

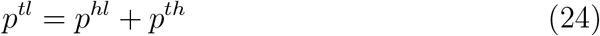

The two capillary pressure-saturation relationships read

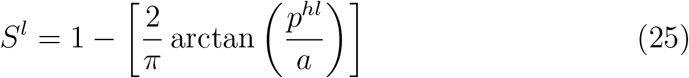

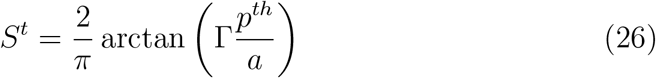

where *a* is a constant parameter depending on the ECM microstructure, and Γ is the ratio of HC-IF and TC-HC interfacial tensions. Parameter *a* was calibrated by *ex vivo* mechanical testing in [41]. In the literature, a generic value for an invasive tumor cell line was previously fixed at 6 [24]. Experimental measurements of surface tension of astrocytes and different glioblastoma cell lines [44, 45] give ratios between 1.3 and 4.8. A higher ratio characterizes a higher invasiveness.

#### Malignant cells mobility

The fraction of GBM cells that expressed a malignant phenotype, *ω*^*ρt*^, influences the dynamic viscosity of the GBM phase, as these cells are more mobile. Beforehand, the dynamic viscosity of the GBM phase is the same as that of healthy glial cells *μ*_*h*_. The influence of the malignant cells fraction follows this equation:

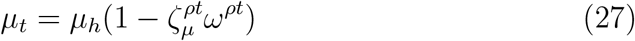

where 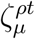 is the coefficient representing the malignant fraction influence, and is to be calibrated.

#### ECM degradation and permeability

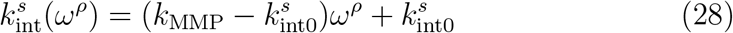

where 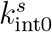 is the intrinsic permeability deduced from imaging data and bounded by experimental literature and *k*_MMP_, corresponding to the permeability of a fully degraded ECM, and is to be calibrated.

#### Oxygen diffusion

The main nutrient considered in our model is oxygen, regulating tumor growth and hypoxia. For the oxygen diffusion, Fick’s law was adapted to a porous medium, to model the diffusive flow of oxygen Eq.10:

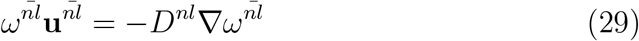

where *D^nl^* the diffusion coefficient for oxygen in the interstitial fluid is defined by the constitutive equation from [43]

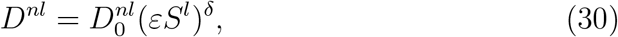

where 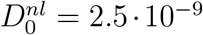 corresponding to the the ideal case of oxygen diffusion in pure water, *i*.*e*. with *εS*^*l*^ = 1, at 37° [46]. The exponent *δ* is equal to 2, to account for the tortuosity of cell-cell interstitium where oxygen diffuse [24].

#### Tumor cells growth and metabolism

Tumor cell growth is related, for its main part, to the exchange allowed by oxygen between the IF and the living fraction of the tumor. For its smaller part, it is related to the exchange allowed by other nutrients (in this case, lipids) in hypoxic situation, between the IF and the positive phenotype IDH fraction of the living tumor. The total mass exchange from IF to the tumor cell phase is defined as

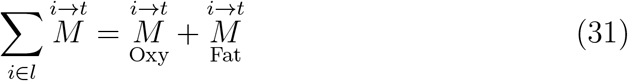

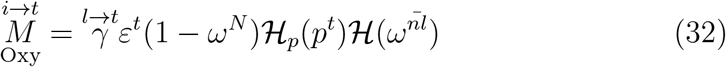

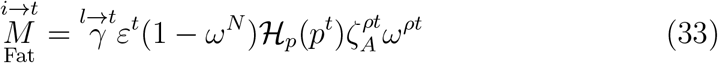

where 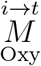 represents the nutrient pathway of TC metabolism and 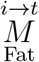 the anoxic growth part due to lipids synthesis of GBM malignant cells [47].

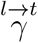 is the tumor growth rate parameter, *ε*^*t*^(1 − *ω*^*N*^) is living fraction of the tumor, *ω*^*ρt*^, its positive IDH phenotype fraction and 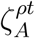, their apoptosis inhibited fraction to calibrate.

ℋ and ℋ_*p*_ are regularized step functions varying between 0 and 1, with two threshold parameters *σ*_1_, *σ*_2_, *i*.*e*. ℋ = ℋ (*σ, σ*_1_, *σ*_2_). When the variable *σ* is greater than *σ*_2_, ℋ is equal to 1, it decreases progressively when the variable is between *σ*_1_ and *σ*_2_ and is equal to zero when the variable is lower than *σ*_1_. ℋ represents the growth dependency on oxygen:

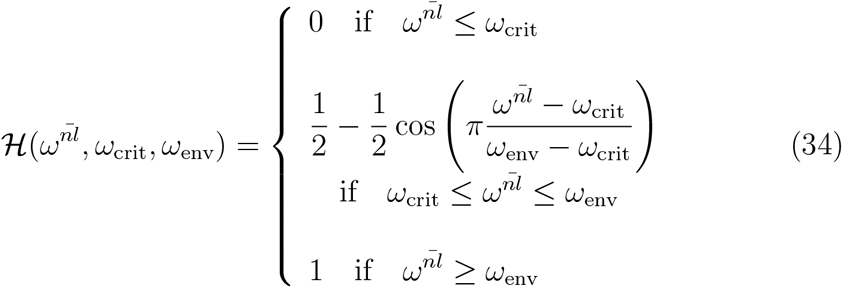

*ω*_env_, the optimal oxygen mass fraction, is set to 4.2·10^−6^ which corresponds, according to Henry’s law, to 90mmHg, the usual oxygen mass fraction in arteries (see [48]). *ω*_crit_, the hypoxia threshold, is cell-line dependent, for tumor cells, it has been set to a very low value: 10^−6^ (≈ 20mmHg, for common human tissue cells, hypoxic level is defined between 10 and 20mmHg [49]).

The function ℋ_*p*_ represents the dependency on pressure:

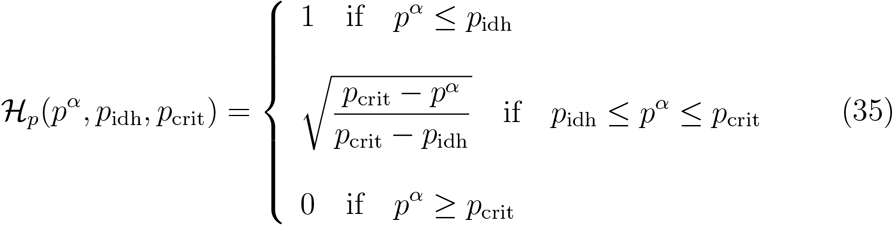

where *p*_idh_ is the minimal threshold of internal pressure that allows GBM cells for switching phenotype and *p*_crit_, the internal pressure threshold which totally stops the GBM cells growth.

Before phenotype switch, IDH wild-type GBM cells are known to produce an important quantity of stroma [29]. Therefore, the fraction 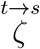 of the mass growth term related to oxygen metabolism 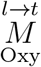 is converted into stroma:

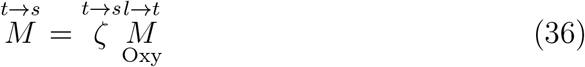

As the tumor grows, oxygen produced by the vascular fraction of the solid scaffold is taken up by the IF phase, giving the following form for the sink and source terms in Eq.10:

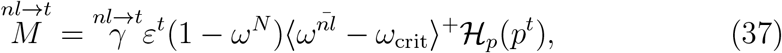

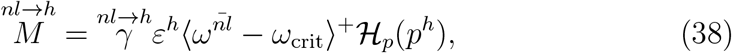

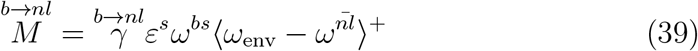

where 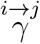 is the corresponding mass exchange rate form phase *i* to phase *j*, where the term *ε*^*t*^(1 − *ω*^*N*^) is the volume fraction of living tumor cells, *ε*^*h*^ the volume fraction of healthy cells and *ε*^*s*^*ω*^*bs*^ the volume fraction of vascularized stroma.

The necrotic growth rate *r*^*Nt*^ is defined by:

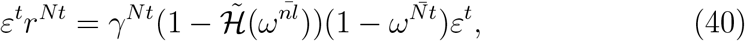

where *γ*^*Nt*^ is the necrotic growth rate. All the parameters to calibrate are summarized Table 3.

### 3. Patient specific image-informed modeling

#### 3.1. Patient dataset

##### The dataset

is composed of MRI methods with a resolution of 256 × 256 × 200. They are displayed Fig.2. The dataset contains:

**Figure 1:**
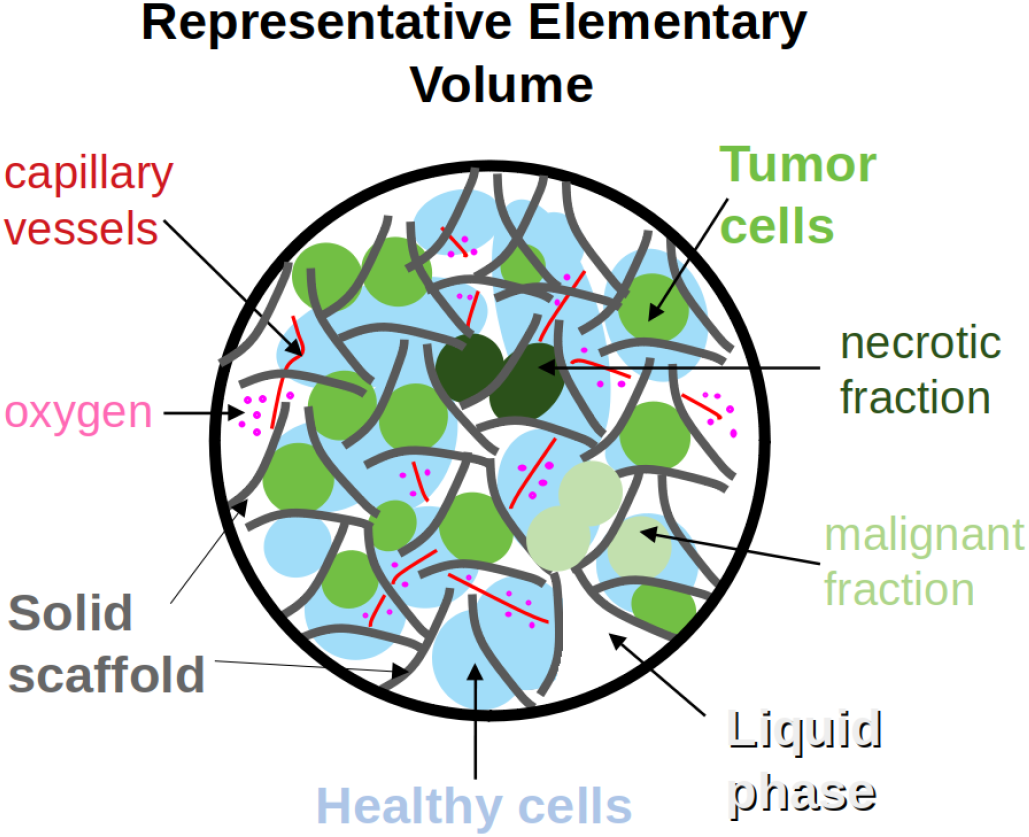
Representative elementary volume of the modeling. Phases are in bold font: solid scaffold (grey), liquid phase (white), healthy cells (blue) tumor cells (green). Species: of the solid scaffold, capillary vessels (red); of the liquid phase, oxygen (pink); of the tumor cells, necrotic (dark green), malignant (light green).

**Figure 2:**
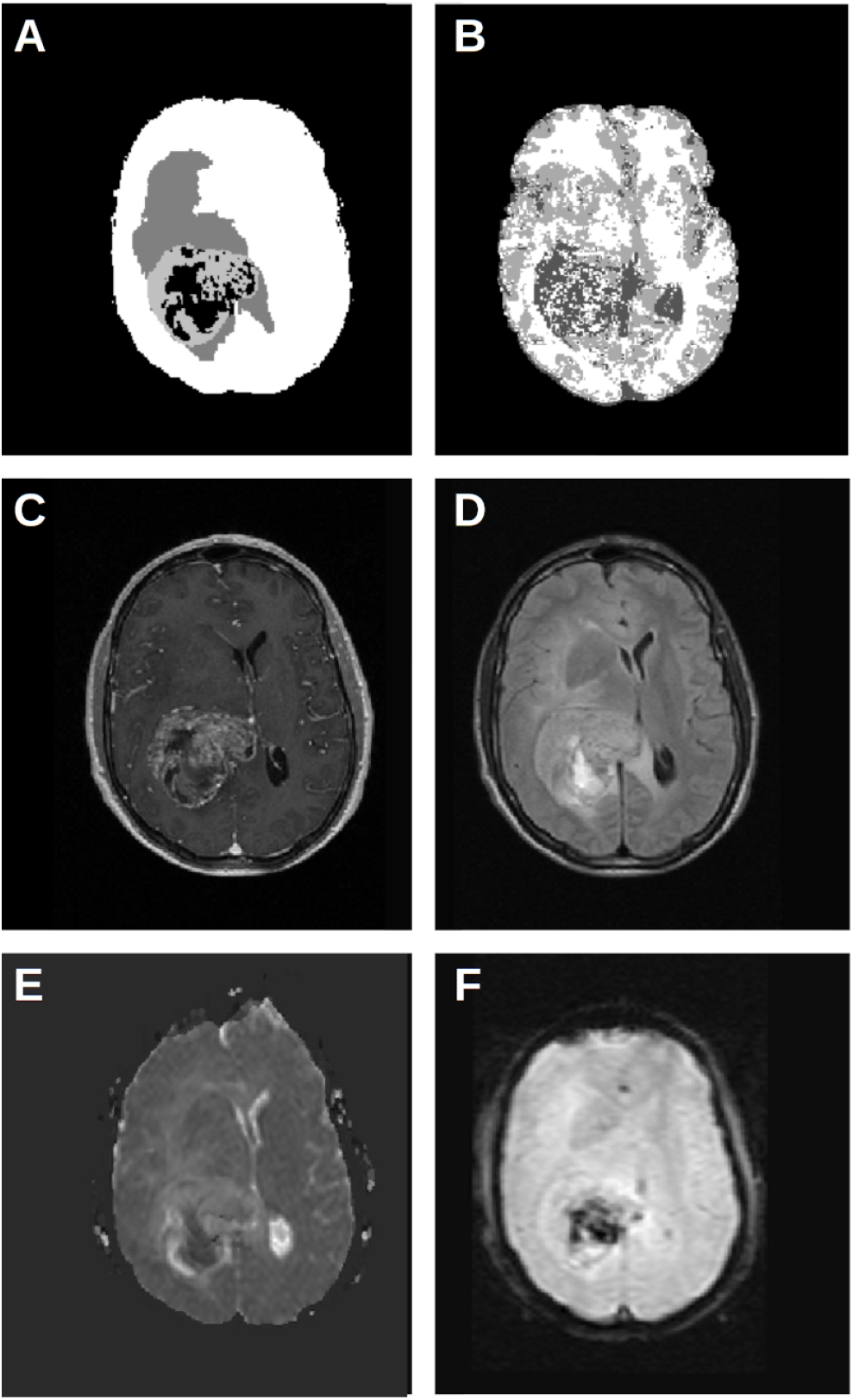
Patient imaging dataset. **A** DeepMedic segmentation gives brain mask (white), edema (dark grey), tumor (light grey) and necrotic (black) zones. **B** FAST segmentation gives only grey and white matter zones, as the tumor tissue is partially misinterpreted as CSF (dark grey). **C** T1-CE method gives the density of solid components (brighter contrast, higher density). **D** FlAIR method gives the density of fluid components (brighter contrast, higher density). **E** ADC method gives diffusion coefficient of water (brighter contrast, higher coefficient). **F** r-CVB method gives the permeability between intra- and extra-vascular space (brighter contrast, higher permeability).

- A segmentation, Fig.2A, by DeepMedic convolutional neural network [50], cleaned by authors of this article. The segmentation gives edema, tumor and necrosis. The segmentation is performed by using the T1 Gadolinium contrast enhanced (T1-CE) method, Fig.2C, and the very long sequence T2 fluid attenuated inversion recovery (FlAIR) method, Fig.2D.
- A segmentation, Fig.2B, by FAST hidden Markov chain [51], which only inform about grey and white matter, as the tumor tissue is partially misinterpreted as CSF. FAST uses T1-CE method for its segmentation.
- A diffusion weighted MRI method, termed as apparent diffusion coefficient (ADC) of water, Fig.2E.
- A perfusion MRI method, termed as relative cerebral blood volume (rCBV), Fig.2F.

This dataset is given at three time points: pre-operative examination, after the 6 cycles of RT-TMZ therapy and after 102 days of TMZ maintenance. 63 days separates the first two time points, and the third time point occurs 165 days after the first one. The first point is used for initial conditions of the model, the second point for the calibration of the parameters and the last for the evaluation of the parameters.

##### Design of the computational domain

The region of interest (ROI) is defined in accordance with surgical practice [52]. The ROI corresponds to the segmented volume of the constrast-enhanced tumor plus 2 cm margin around this volume, where a GBM has the greater probability to progress. The computational domain is shown Fig.3B. It contains the ROI and an additional margin between 4 and 7 mm. This additional margin was designed to ensure that the prescribed boundary conditions on the computational domain have a negligible influence on the GBM evolution. The boundary conditions can be found Section 3.3 and the design process is described in details Appendix Appendix A.

**Figure 3:**
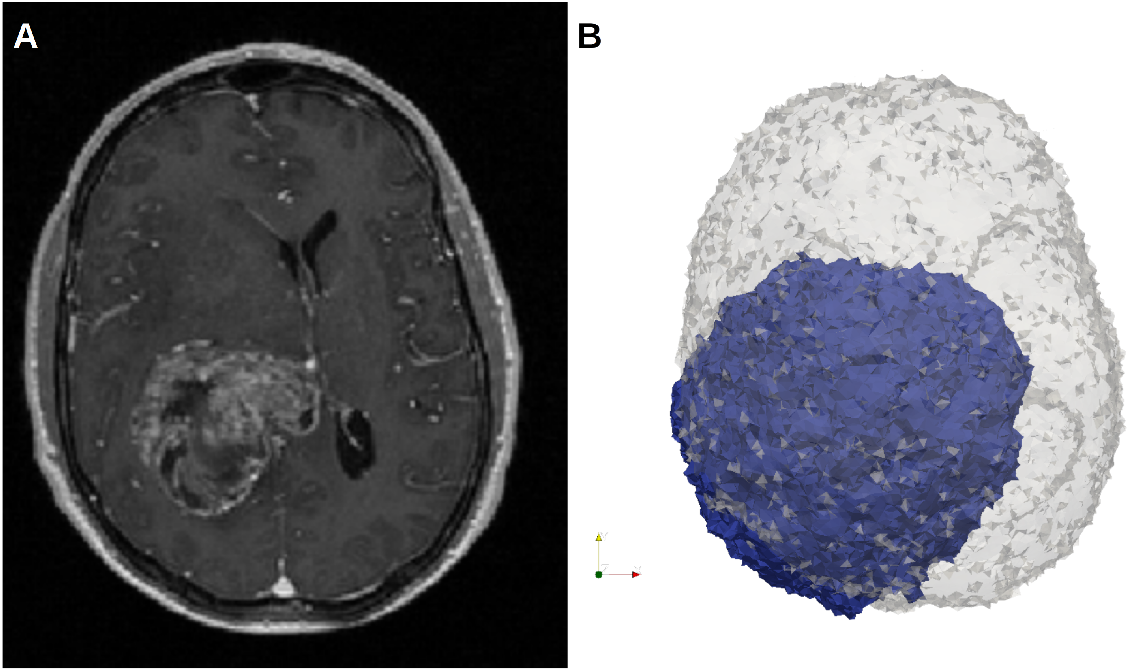
Design of the computational domain. **A** Patient data, axial view of T1-CE MRI method. **B** 3D view; Brain mask (transparent grey) extracted from patient data T1-CE; Computational domain (blue) defined by boundary conditions 2.27 ± 0.3 cm around the tumor zone. These conditions have a negligible influence on the tumor evolution, for further details see section 3.3 and appendix Appendix A.

#### 3.2. Initial parameters settings

The quantities and methods are summarized Tables 1, 2, 3 and 4. These tables summarize 1) the parameters obtained by *ex vivo* mechanical testing, 2) the parameters informed by clinical imaging, 3) the parameters that require calibration and 4) the treatment parameters.

**Table 1:**
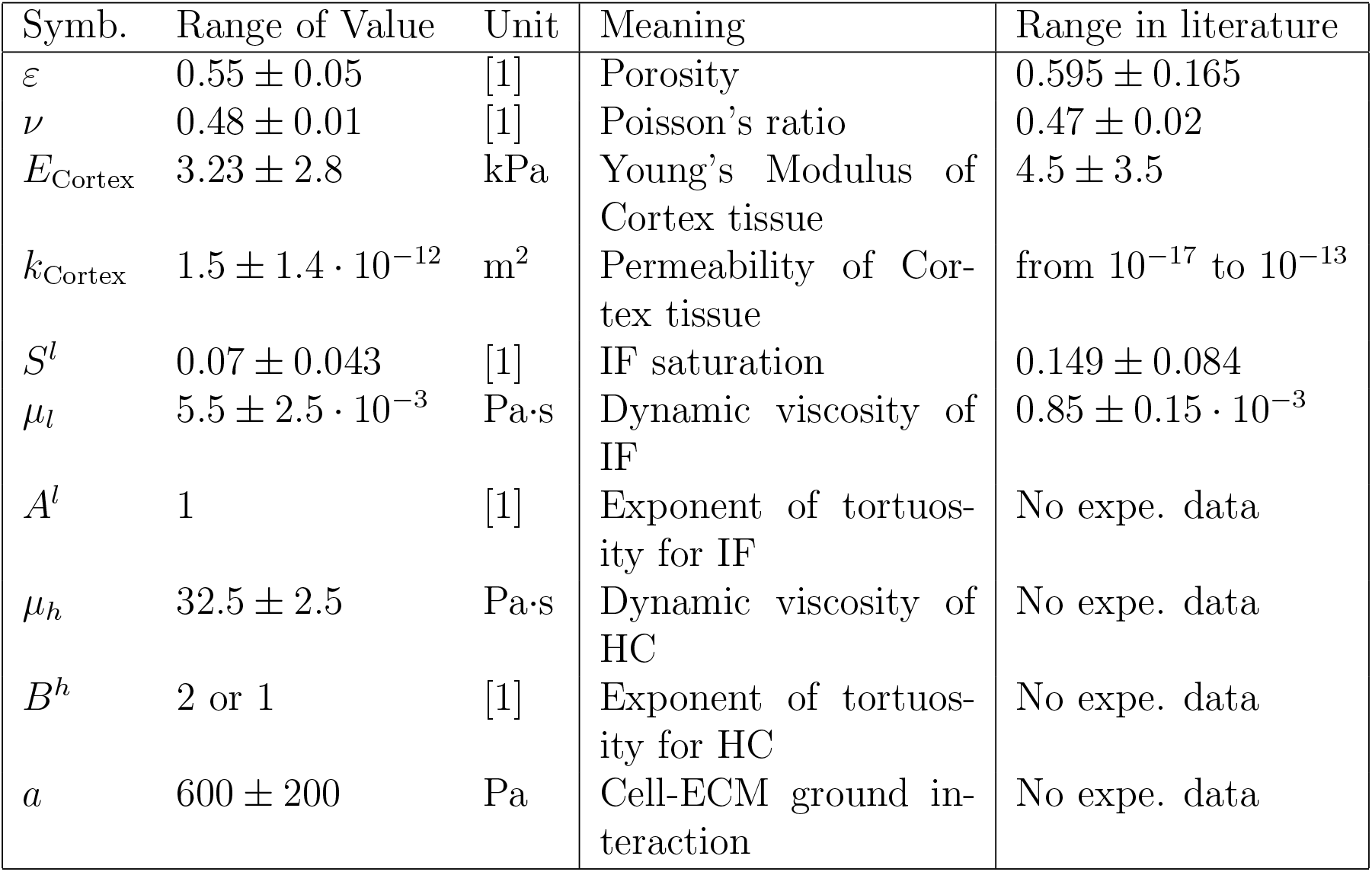
Parameters estimation by *ex vivo* mechanical testing [41]. For the sources of the literature values, see [41].

**Table 2:** Parameters deduced from MRI methods.

| Type | Symb. | Unit | Method(s) | Parameter |
| --- | --- | --- | --- | --- |
| **Material parameters** | $\bar{k}()$ | m<sup>2</sup> | ADC, Segment.[51] | Permeability mapping |
| | $E()$ | Pa | Segment.[50] | Young's Modulus mapping |
| **Initial conditions** | $p^l$ | Pa | ADC | Interstitial fluid pressure |
| | $p^{hl}$ | Pa | ADC, Segment.[50] | Healthy cells pressure |
| | $p^{th}$ | Pa | ADC, Segment.[50] | Tumor cells pressure |
| | $\omega^{bs}$ | [1] | rCVB | Vascular fraction |

**Table 3:** Model’s parameters to be calibrated, initial values, and sources.

| Type | Symb. | Value | Unit | Meaning | Source |
| --- | --- | --- | --- | --- | --- |
| **Poromechanics** | $E_{\text{IDH}}$ | 4000 | Pa | Young's Modulus of Cross-linked ECM | interpreted from [41] |
| | $p_{\text{crit}}$ | 1500 | Pa | Critical threshold of mechanical inhibition | - |
| | $\Gamma$ | 6 | [1] | Interfacial tension ratio between HC-IF and TC-HC | [24] |
| | $k_{\text{MMP}}$ | $10^{-10}$ | $\text{m}^2$ | Maximum permeability of the ECM degraded by MMP | - |
| **Oxygen biology** | $\overset{l \rightarrow t}{\gamma}$ | $2.16 \cdot 10^{-2}$ | $\text{kg}/(\text{m}^3 \cdot \text{s})$ | TC growth rate | - |
| | $\overset{nl \rightarrow t}{\gamma}$ | 3.5 | $\text{s}^{-1}$ | TC oxygen consumption rate | - |
| | $\overset{nl \rightarrow h}{\gamma}$ | $2.5 \cdot 10^{-1}$ | $\text{s}^{-1}$ | HC oxygen consumption rate | - |
| | $\overset{b \rightarrow nl}{\gamma}$ | $1.44 \cdot 10^{-2}$ | $\text{s}^{-1}$ | Capillaries oxygen production rate | - |
| | $\omega_{\text{crit}}$ | $8 \cdot 10^{-7}$ | $\text{kg}/\text{m}^3$ | GBM cells, hypoxic threshold oxygen mass fraction | interpreted from [49] |
| | $\gamma^{Nt}$ | $10^{-2}$ | $\text{kg}/\text{m}^3$ | Necrotic core growth rate | [24] |
| | $\omega_{\text{crit}}^h$ | $10^{-6}$ | $\text{kg}/\text{m}^3$ | Glial cells, hypoxic threshold oxygen mass fraction | [49] |
| **ECM mechano-biology** | $p_{\text{idh}}$ | 1000 | Pa | Phenotype switch mechanical threshold | - |
| | $\zeta_A^{\rho t}$ | $1 \cdot 10^{-1}$ | [1] | TC apoptosis inhibited fraction | - |
| | $\zeta_\mu^{\rho t}$ | $9.9 \cdot 10^{-1}$ | [1] | TC dynamic viscosity loss fraction | - |
| | $\overset{t \rightarrow s}{\zeta}$ | $4 \cdot 10^{-1}$ | [1] | Stroma production coefficient | - |
| | $\zeta^\rho$ | 2.5 | [1] | Phenotype switch coefficient | - |

**Table 4:**
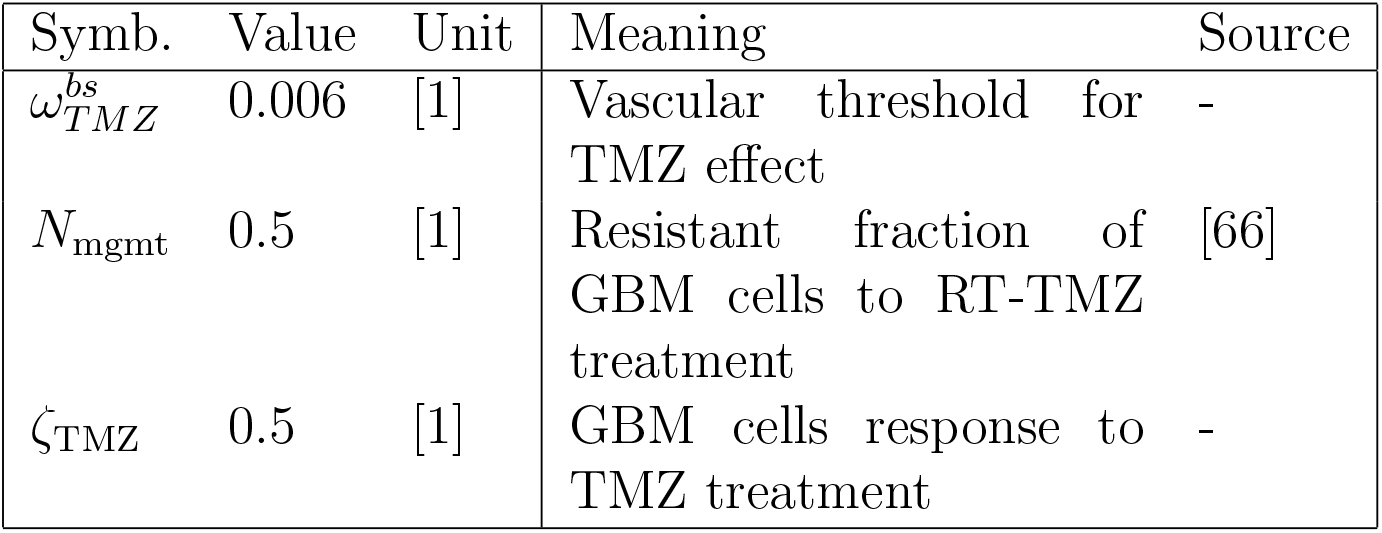
RT-TMZ treatment parameters.

##### Segmentation

Tumor segmentation with Deep Medic and brain segmentation with FAST give two distinct partitions of the computational domain Ω:

- Deep Medic partition gives Ω_*CE*_, the GBM Contrast Enhanced domain, Ω_*N*_, the GBM necrotic domain, Ω_*E*_ the GBM edema domain and Ω_*O*_ the outer segmentation domain. The Deep Medic partition is defined by Ω_*CE*_ ∪ Ω_*N*_ ∪ Ω_*E*_ ∪ Ω_*O*_ = Ω.
- FAST partition gives Ω_*CSF*_, the CSF compartment of the patient brain -which cannot be exploited due to tumor tissue-, Ω_*G*_, the grey matter subdomain and Ω_*W*_ the white matter subdomain. The FAST partition is defined by Ω_*CSF*_ ∪ Ω_*G*_ ∪ Ω_*W*_ = Ω.

##### ADC method

gives diffusion the coefficient of water which is inversely correlated to the cell density [17]. For this reason, we choose to consider the interstitial fluid (IF) saturation *S*^*l*^ as proportional to the ADC contrast. As there is presence of an edema, the maximum contrast ADC_max_ corresponds to a pathological value of IF pressure. We set it to *p*_max_ = 400Pa, which corresponds to a 3 mmHg increase in the intracranial pressure defined in Eq.13. The minimal value (ADC_min_), which corresponds to the maximum cellularity (*e*.*g*., the tumor necrotic zones), is set at a value below normal pressure *p*_min_ = 40Pa (+0.29 mmHg). Then, we obtain the linear function to prescribe the initial conditions for IF pressure:

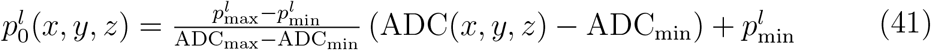

The tumor quantities are defined over Ω_*CE*_ ∪ Ω_*N*_ and they both need segmentation and MRI methods to be specified. In [53], histological cuts on 7 patients with GBM gives a volume fraction of GBM cells of *εS*^*t*^ = 0.12 ± 0.07. With the porosity estimation in [41], *ε* = 0.55 ± 0.05, we obtain a range of tumor cells saturation *S*^*t*^ between 0.115 and 0.24. As base values, we chose the maximum saturation in the necrotic core 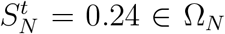 and a value slightly below average in the contrast-enhanced zone 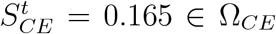. The relationship between *S*^*t*^ and TC pressure difference *p*^*th*^ depends on two parameters *a* and Γ. With *a* = 550 and Γ = 6, we obtain 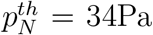 and 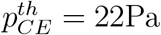. These base values are tuned by the mean of the ADC mapping:

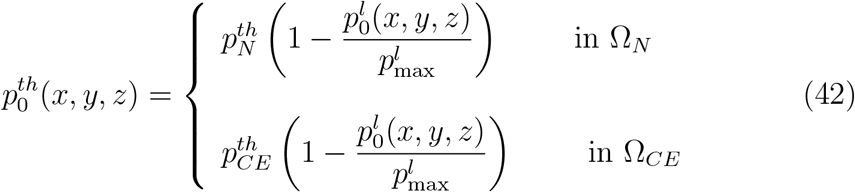

As the saturation of healthy glial cells *S*^*h*^ is constrained by Eq.2, this saturation is not directly linked to the capillary pressure *p*^*hl*^ of glial cells. Nevertheless, the IF saturation *S*^*l*^ is subjected to *p*^*hl*^ by the Eq.25. The initial mapping of *p*^*hl*^ with DeepMedic segmentation and the ADC method respects the range of physical values deduced from [41]. In the edema zone Ω_*E*_, the base value of 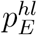 is fixed at 800 Pa, which gives a higher value of IF *S*^*l*^ = 0.39, which is pathological. In the rest of the domain Ω \ Ω_*E*_, the base value 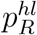 is fixed at 1.6 kPa, which gives a physiological value of *S*^*l*^ = 0.11.

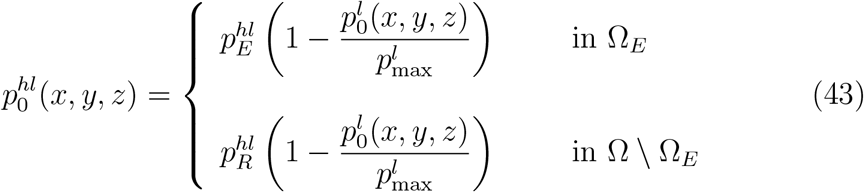

The initial value of *p*^*hl*^ also sustains the intracranial pressure *p*^*s*^ (see Eq.13) which is defined all over the domain. With the initial mapping of *p*^*hl*^, we obtain an average intracranial pressure *p*^*s*^ between 7.75 mmHg and 12.5 mmHg. These values are in balance with physiological measurements [54, 55]. Therefore, we choose the initial values of *p*_idh_ and *p*_crit_ within this range: *p*_idh_ is set to 1 kPa (≈ 7.5 mmHg) and *p*_crit_ is set to 1.5 kPa (≈ 11 mmHg).

The mapping of the intrinsic permeability of the stroma 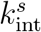 is performed through FAST segmentation, as the white matter tracts have a higher permeability [56], and through the ADC method, because we interpret the zones of accumulation of fluid as zones with a higher permeability. As the patient data does not contain a diffusion tensor imaging method, 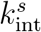 remains a scalar heterogeneous quantity, not a vectorial quantity. The determination of intrinsic permeability of the brain is a very difficult experimental task, and the wide range of values obtained remains an open debate (see [41] for details). For 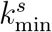, we choose a one order lower value than we found in [41], 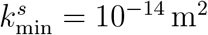.

For 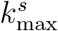, we choose the lower bound of [41], 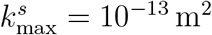, as the grey and white matter were not distinguished. Where the voxels are labelled as white matter, we follow the trends of the results of Jamal *et al*. [56] by prescribing a 15 fold value for 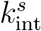: its maximum value of is 1.5 · 10^−12^ m^2^.

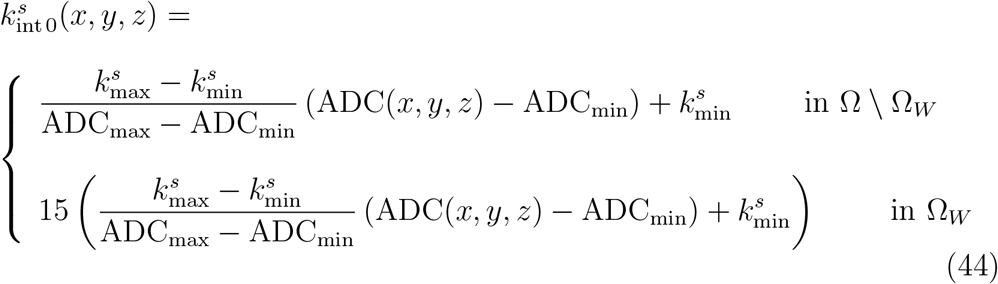

##### rCBV method

is treated the same way as the ADC method. It provides the vascular fraction *ω*^*bs*^ of the solid scaffold *ε*^*s*^. The maximum contrast rCBV_max_, which corresponds to a neo-vascular network, sets the vascular fraction to 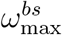. From early work on angiogenesis, Folkman *et al*. in [57] estimated the vascularized fraction of a subcutaneous tissue undergoing angiogenesis to 1.5%, which is 400 times that of healthy tissue. However, cortex tissue is already a highly vascularized tissue, with a volume fraction estimated to be between 3% and 5% (see Yiming *et al*. in [58]). We chose to set 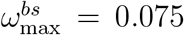, 50% higher than maximal physiological value, and 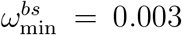, 10 times lower than minimal healthy value for poorly vascularized zones. We obtain for the vascular fraction of the solid scaffold:

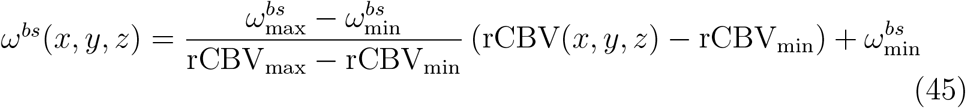

##### General mechanical parameters of cortex tissue

were prescribed our previous work [41]. In [41], we reproduced two mechanical tests on healthy human and animal cortex: confined compression (*N* = 6), *i*.*e*. consolidation tests of Franceschini *et al*. [59] and unconfined compression (*N* = 40), *i*.*e*. indentation tests with several load rates and diameters of Budday *et al*. [60]. Part of the experimental results were used for calibration and another part for validation (*N* = 3). All the details are provided in [41]. This article allows for reducing the range the mechanical parameters of cortex tissue provided in the literature [61, 62, 63], and more specifically in the poromechanical literature [64, 56]. Although individual variation could be considered, these parameters are related the general mechanical behavior of healthy tissue. They are presented in Table 1. These parameters will be thereafter considered as fixed.

##### Partial resolution of the mathematical system

Imaging data or mechanical tests do not give information on the initial state of the oxygen fraction 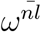. Physiology literature gives information on the bounds of the tumor growth metabolism (Eq.34). The hypoxia threshold *ω*_crit_, in a brain tumor environment, is estimated to an oxygen fraction between 4.5 · 10^−7^ and 10^−6^ [49]. These values correspond, according to Henry’s law, to an oxygen partial pressure between 10 mmHg and 22.5 mmHg. The clinical measurements in [65] give a range between 30 and 48 mmHg for physiological values in brain tissue. Therefore, we set the oxygen fraction of healthy brain tissue *ω*_env_ to 1.9 10^−6^, which corresponds, according to Henry’s law, to 42.5 mmHg. Nevertheless, between the two bounds defined by *ω*_crit_ and *ω*_env_, the oxygen fraction at each voxel of the domain is not known. To fix this, the mathematical system is partially solved. The oxygen fraction is set to 10^−6^ in Ω_*CE*_ ∪ Ω _*N*_ and to 1.9 · 10^−6^ in the remaining part. With these initial conditions, the system is solved with a very small time increment (*dt* = 1 s), as this initial state is very unstable. When the system becomes steady, *i*.*e*. the variation of the oxygen fraction in one second becomes negligible, the simulation is stopped and the solution of 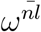 is stored as initial condition for this unknown. This computation corresponds to 90 seconds of simulated time.

The same situation occurs for the displacement field **u**^**s**^, as the previous deformation of the organ is not recorded. As the fluid phases exert a pressure on the stroma, the same process than for 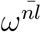 is adopted. When a mechanical steady state is reach between fluid pressures and stroma displacement, the simulation is stopped and the resulting displacement field **u**^**s**^ is conserved as the initial solution for this vectorial unknown. The corresponds to 6 minutes of simulated time, the initial **u**^**s**^ reaches a maximum displacement between 50 *μ*m and 60 *μ*m.

##### Clinical literature

Kitange *et al*. in [66], showed by *in vitro* experiments and animal models that MGMT activity greatly increase the GBM resistance to TMZ treatment, whereas the methylation of MGMT decrease its activity and allows for a better response. Their statistical analysis of *in vitro* results showed an increase between 50% and 60% of the surviving GBM cells fraction with a non-methylated (n-)MGMT marker. Based on these findings, we set the resistant fraction of n-MGMT marker to 0.5 as initial guess in Eq.18. Without clinical measurement, the minimum vascular fraction required to convey the effect of TMZ is set to 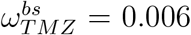, this value corresponds to twice the minimal vascular fraction. These parameters are summarized in Table 4.

#### 3.3. In silico reproduction process

##### Finite element formulation

We implemented the above model with Dolfin, the C++ libraries of the FEniCS framework [67, 68]. We used an incremental formulation, *i*.*e*. 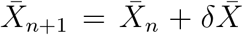, for the mixed finite element (FE) formulation. We resolve the system by the means of a fixed-stress staggered scheme: the pressures are solved with a fixed stress tensor, the stress tensor is solved with the updated pressures, and the loop is subjected to the norm of the solution increment as convergence criterion (for instance, see [69]). All the codes are available as supplementary material and can be downloaded at https://github.com/SUrcun/GBM_mecano_bio.

##### Boundary conditions

All unknowns are subjected to a homogeneous Dirichlet conditions on the domain boundary. This is a consequence of the incremental formulation. For each unknown *α*, we prescribed *δX*_*α*_ = 0 on ∂Ω, the boundary of the domain. In other words, the initial settings of the unknowns remain unchanged at the boundary of the domain during the simulation. The influence of the boundary distance on the FE solution is studied in Appendix Appendix A.

##### Quantities evaluated

In a multiphase system, grasp the relevant quantities is not always straightforward, for instance, the saturation of tumor cells *S*^*t*^ could be meaningless without the indication of the porosity *ε*. For instance, if we want to delineate a tumor area, the significance of a high *S*^*t*^ could be diminished by a small *ε*. Hence, we adopt the following measure for the interpretation of the results:

The volume fraction of tumor cells:

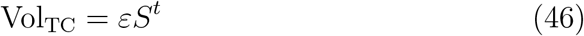

Vol_TC_ can be separate in three relevant quantities, the living tumor cells:

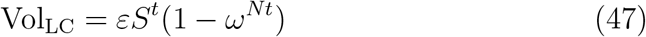

The malignant tumor cells:

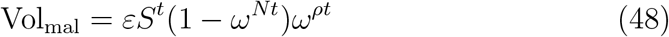

The necrotic tumor cells:

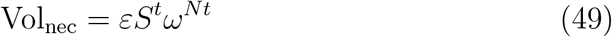

##### Error measure

To measure the quality of the numerical results, we followed the prescription of [70]: the root mean square error (RMSE) relative to a reference, which is specified accordingly. The RMSE of the numerical quantity *ξ*_num_ relative to the patient reference *ξ*_data_, evaluated at *n* points is computed as:

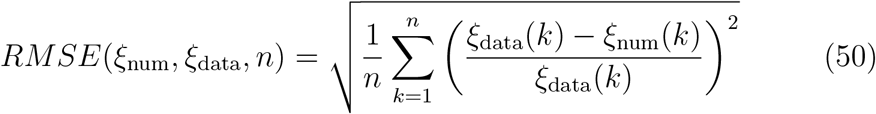

##### Sensitivity analysis

*cost functions and Sobol indices ..* We performed a local sensitivity analysis to estimate Sobol indices on the patient calibration dataset, to assess the sensitivity of the computational outputs to the input parameters. First, we designed the cost function *J*_over_, which quantify the error between the numerical results and the patient calibration dataset, by measuring the spatial overlapping:

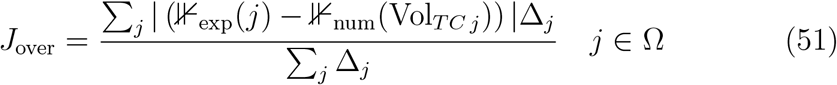

with

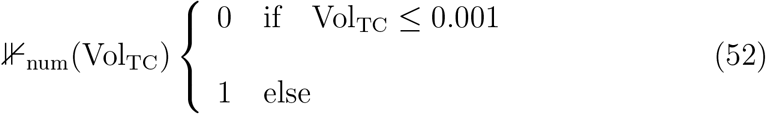

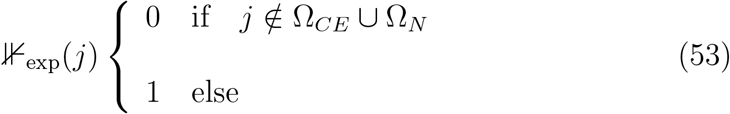

where Δ_*i*_ is the volume of the *i*^*th*^ tetrahedra, where 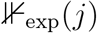 is the characteristic function of the patient segmentation at the second time point - *i*.*e*. the calibration dataset -, and 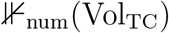 the characteristic function of the computational GBM at the same time point.

The 16 parameters at their initial values (see Table 3) give 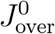. Then, the 16 parameters are perturbed one at a time, in four configurations: ±10% and ±20%. The cost variation 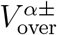 of a parameter α is defined by:

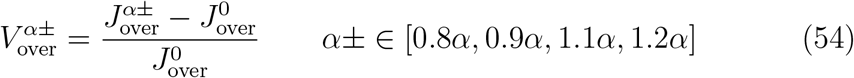

Then, the points of the variation are linearly interpolated. The influence of the parameter *α* is deduced from the slope *θ*_*α*_ of the linear fit. The first-order Sobol index *S*_*α*_ is calculated as follows:

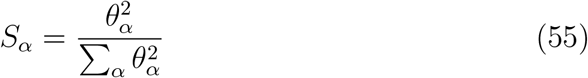

##### Calibration

To minimize *J*_over_, we chose the subset of parameters *α*_*i*_ that gather 90% of the variance, *i*.*e*. 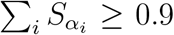. This parameters set is then calibrated using a Newton-Raphson algorithm.

#### 4. Results

##### Local sensitivity analysis

The results of the first order sensitivity analysis are shown in Fig.4, and the values of the Sobol indices in Table 5. The results give that only 6 parameters gather 93.8% of the variance. The hypoxia threshold the GBM cells *ω*_crit_ (Sobol indice 0.213), the interfacial tension between GBM cells and its surrounding medium Γ (0.189), the mechanical pressure required for the phenotype switch *p*_idh_ (0.167), the oxygen production by micro-capillaries 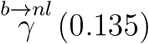, the GBM cells growth rate 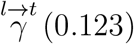 and the oxygen consumption of GBM cells 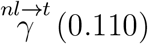. This subset of 6 parameters is calibrated, and the others are considered fixed.

**Figure 4:**
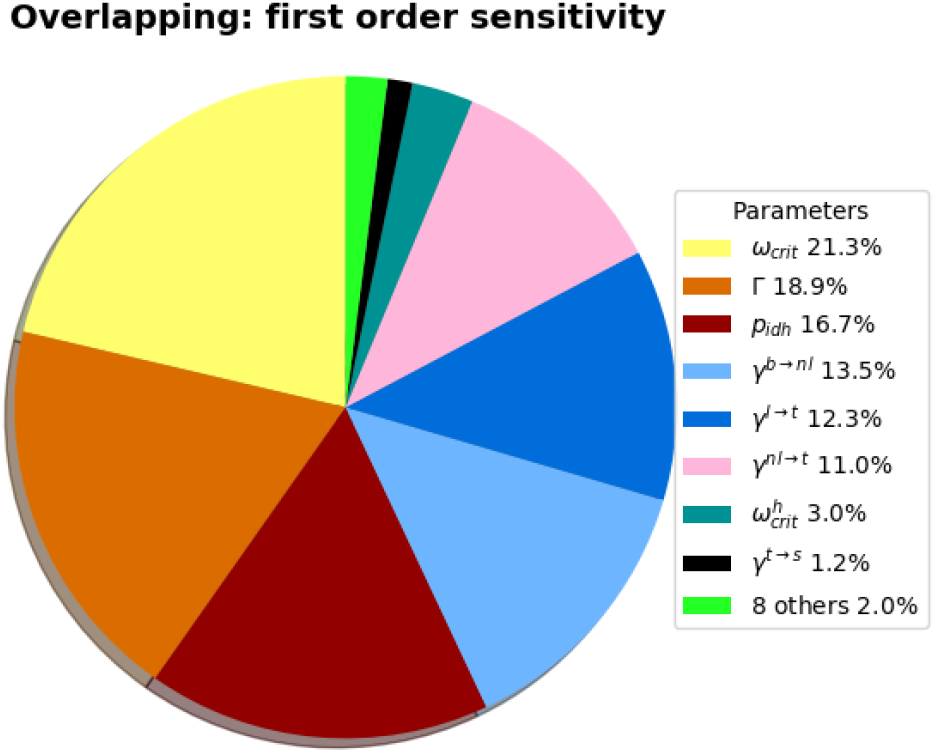
Sobol indices of the parameters at their initial values. Details of the parameters Table 5. 6 parameters gather 93.8% of the variance: *ω*_crit_ (0.213), Γ (0.189), *p*_idh_ (0.167), 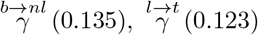 and 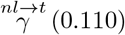. They constitute the parameters subset to calibrate.

**Table 5:**
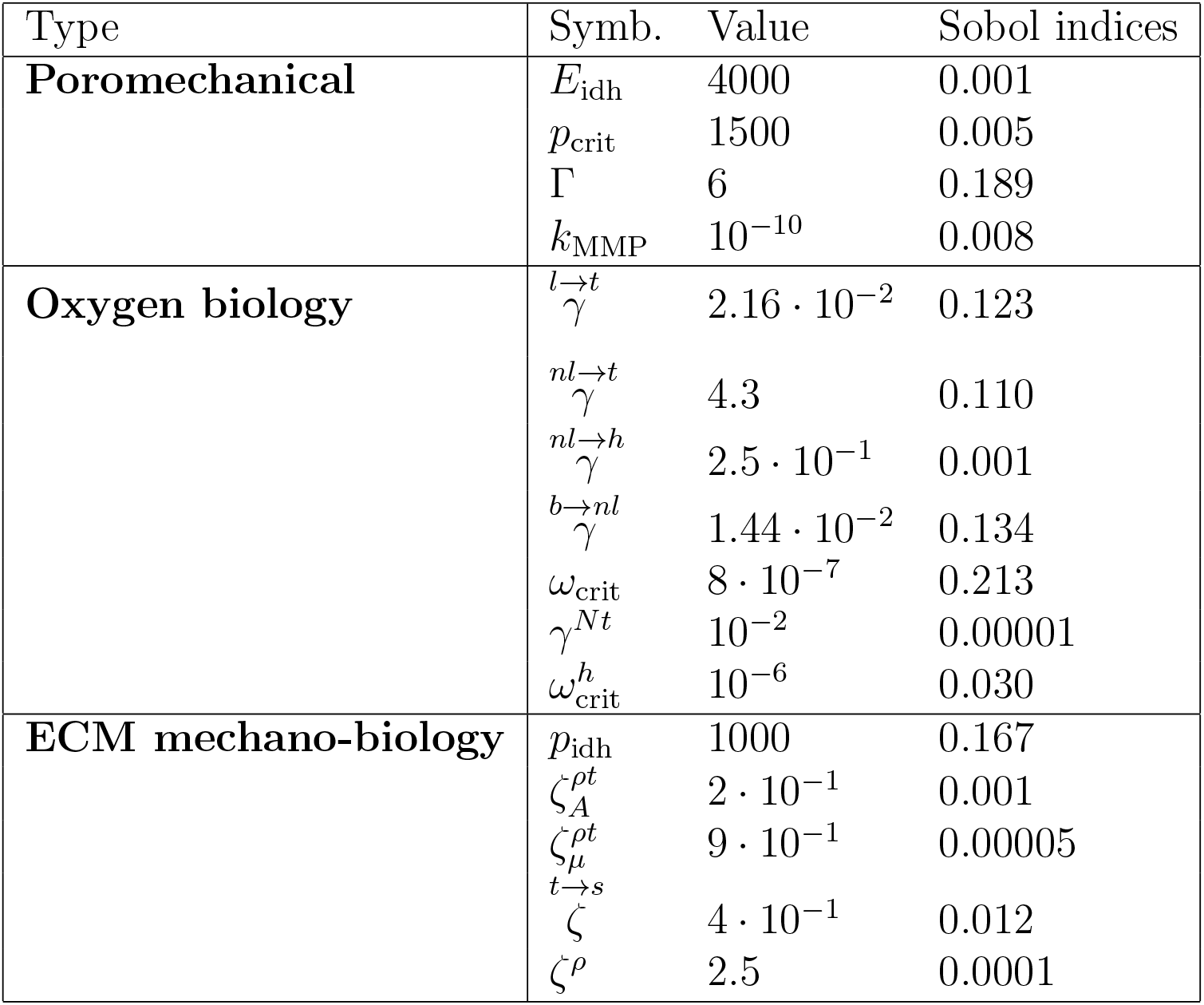
Sobol indices of the parameters at their initial values, perturbed one at a time, in four configurations: ±10% and ±20%. *J*_over_ = 0.581

##### Calibration

The 6 parameters are identified with a Newton-Raphson algorithm, only 5 iterations were performed - with a duration of 4 days of computational time per iteration - giving the following overlapping errors between numerical results and patient dataset at *T*_0_ + 63 days: 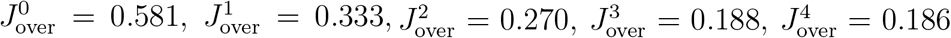 and 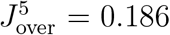. The calibrated values are given Table 6. The 3D results at 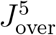 are shown Fig.5. At the fifth iteration, the volume of the simulated tumor is 116.3 cm^3^, the volume of the patient tumor at *T*_0_ + 63 days being 122.5 cm^3^. Then, we obtained a tumor with 5.0% of error in volume and which overlaps 81.4% of the patient tumor.

**Figure 5:**
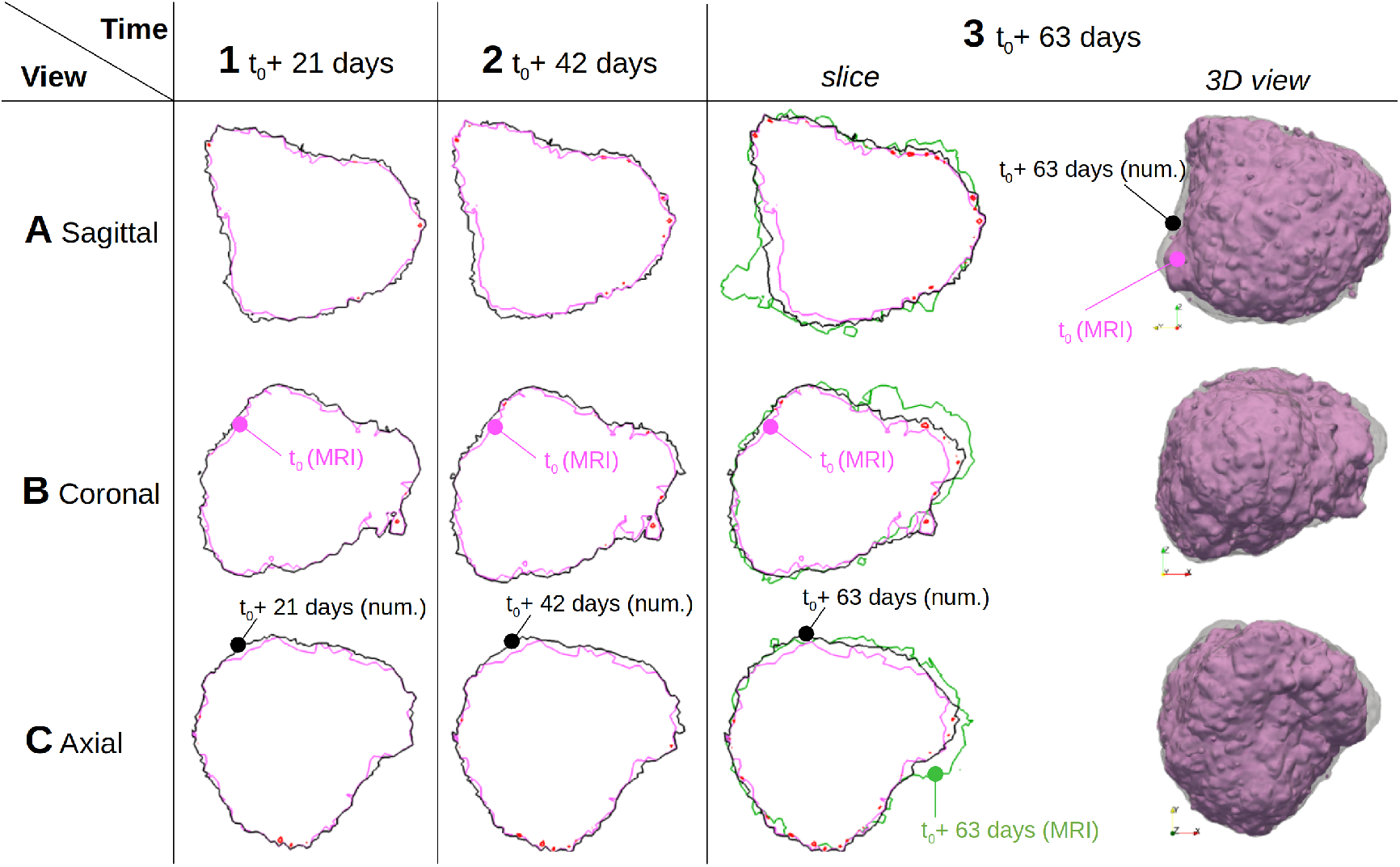
Influence of mechanical inhibition of tumor growth. **A** Sagittal view (along *x* axis). **B** Coronal view (along *y* axis). **C** Axial view (along *z* axis).**0**. **1, 2, 3** slices centered at *x* = −0.0180, *y* = −0.0104, *z* = 0.0416; isoline of patient data Vol_TC_ = 0.001 at *T*_0_ (purple), isoline of inhibiting pressure *p*_crit_ (red). **1** at *T*_0_ + 21 days, isoline Vol_TC_ = 0.001 (black). **2** at *T*_0_ + 42 days, isoline Vol_TC_ = 0.001 (black) after 3 cycles of RT-TMZ treatment. **3** at *T*_0_+63 days, isoline Vol_TC_ = 0.001 (black); isoline of patient data Vol_TC_ = 0.001 (green) after 6 cycles of RT-TMZ treatment. 3D isosurface Vol_TC_ = 0.001 of model outputs at *T*_0_ (purple) and at *T*_0_ +63 days (grey transparent). In the zones where the inhibiting pressure *p*_crit_ is reached, the numerical results show almost no progression. The few zones without this inhibiting pressure correspond to the major progression zones in the patient data. Detail of the mechanism underlying these numerical progression are given Fig.6.

**Table 6:** Parameters calibration, *J*_over_ = 0.186.

| Symb. | Value |
| --- | --- |
| $\omega_{\text{crit}}$ | $7 \cdot 10^{-7}$ |
| $\Gamma$ | 4.5 |
| $p_{\text{idh}}$ | 910 |
| $b \rightarrow nl$<br>$\gamma$ | $1.62 \cdot 10^{-2}$ |
| $l \rightarrow t$<br>$\gamma$ | $2.6 \cdot 10^{-2}$ |
| $nl \rightarrow t$<br>$\gamma$ | 4.1 |

##### Qualitative results

In this section, we define three types of GBM evolution: inhibited, pro-liferative, invasive. Examples of these zones are shown Fig.6. We termed inhibited a zone that reaches the critical threshold *p*_crit_ which mechanically impedes the cell growth. Conversely, proliferative zones do not undergo this mechanical pressure and the cells grow normally. Invasive zones indicate a zone of progression despite a mechanical impediment, *i*.*e*. the GBM progresses in this zone by invasion and not by proliferation.

**Figure 6:**
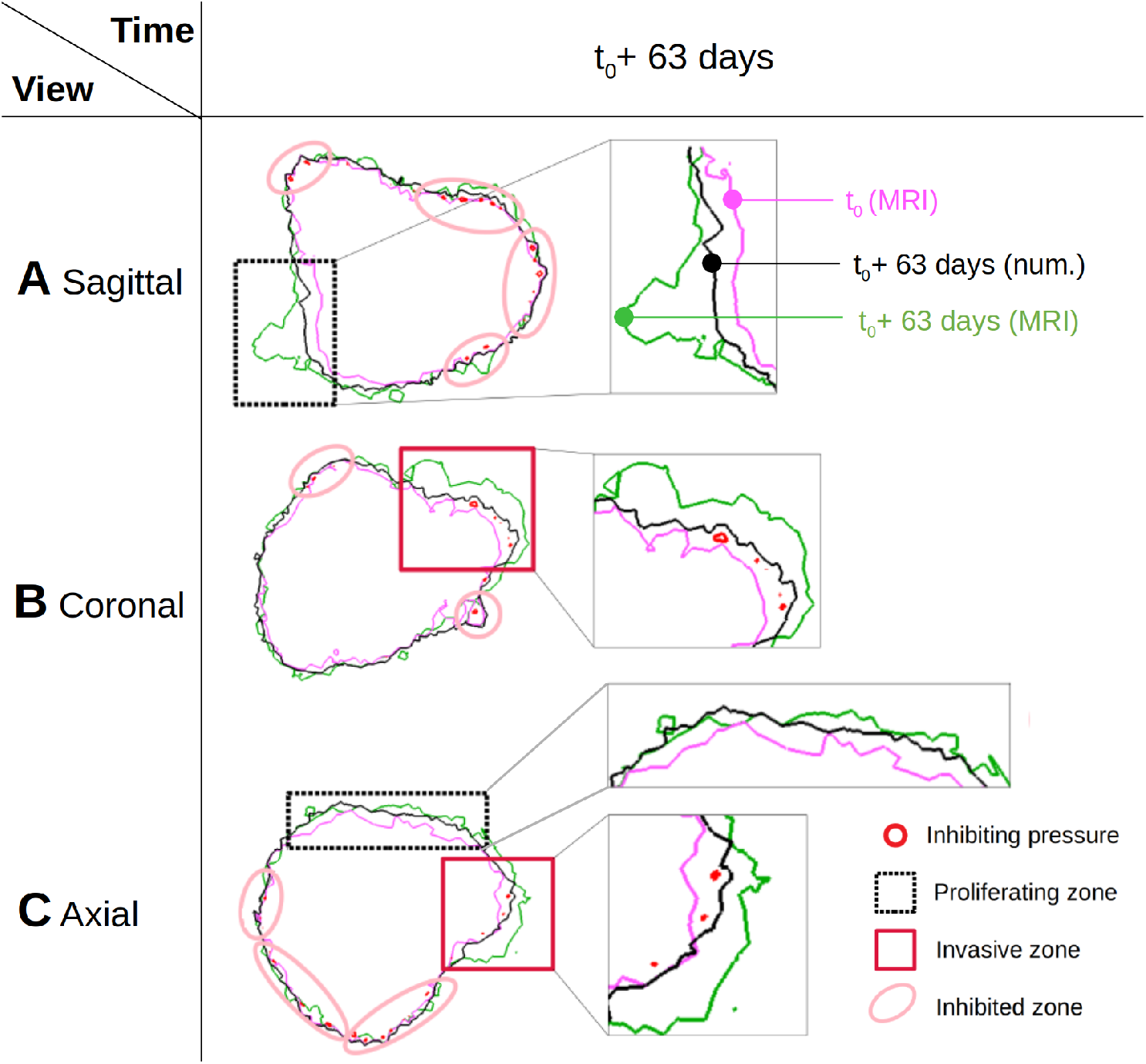
Influence of mechanical inhibition of tumor growth, focus. **A** Sagittal view (along *x* axis). **B** Coronal view (along *y* axis). **C** Axial view (along *z* axis). Isoline Vol_TC_ = 0.001 at *T*_0_ (purple). At *T*_0_ + 63 days after 6 cycles of RT-TMZ treatment: isoline Vol_TC_ = 0.001 (black); isoline of patient data Vol_TC_ = 0.001 (green); isoline of inhibiting pressure *p*_crit_ (red); proliferating zone (black dotted square); invasive zone (red square); inhibited zone (pink ellipse). Almost all inhibited zones correspond to patient data with no growth. Proliferating zones correspond to a TC growth which is not impeded by mechanical pressure, 2 zones of progression in the patient data correspond to this definition in the numerical results (one in sagittal view, one in axial view). Invasive zones correspond to TC migration, as the proliferation is impeded by mechanical pressure, 2 zones of progression in the patient data correspond to this definition in the numerical results (one in coronal view, one in axial view). This migration is explained in detail Fig.7 and 8.

##### Mechanical inhibition of tumor growth

This phenomenon is well documented *in vitro* [71, 72, *73], in vivo* [74], and already used in image-informed model for breast [18] or prostate [19] cancers and were comprehensively reviewed by Jain *et al*. in [75] and more recently by Nia *et al*. [76]. Even if each cell line has its own conditions (inhibiting pressure threshold, shear stress dependency, phenotype switch window, coupling phenomena with hypoxia), the mechanical inhibition of tumor growth is now accepted as a phenomenon shared by many cancers. Specifically for GBM mechanical growth inhibition, to our knowledge, we found only one quantitative study of Kalli *et al*. [77], which estimates for the GBM A172 cell line an inhibiting threshold around 3.5 kPa. In the present study, the inhibiting threshold *p*_crit_ was set, as initial guess, to 1.5 kPa (≈ 11mmHg) but this parameter is not calibrated because its weight in the sensitivity study was to low (Sobol index 0.5%). We see in Fig.5A3 and C3 that several large zones with no progression in the patient data. Almost all these zones correspond, in the numerical results, to zones which undergo a pressure at least equal to *p*_crit_, termed inhibited zones. However, few zones of progression in the patient data (see Fig.5B3) correspond to numerical progression despite the inhibiting pressure. These zones are termed invasive zones. The details on each of these zones is provided in Fig.6. If proliferative zones correspond to a TC growth which is not impeded by mechanical pressure, invasive zones correspond to TC migration after a phenotype switch, as their proliferation is impeded by mechanical pressure. The mechanisms of this migration are explained in detail next paragraph and Fig.7,8.

**Figure 7:**
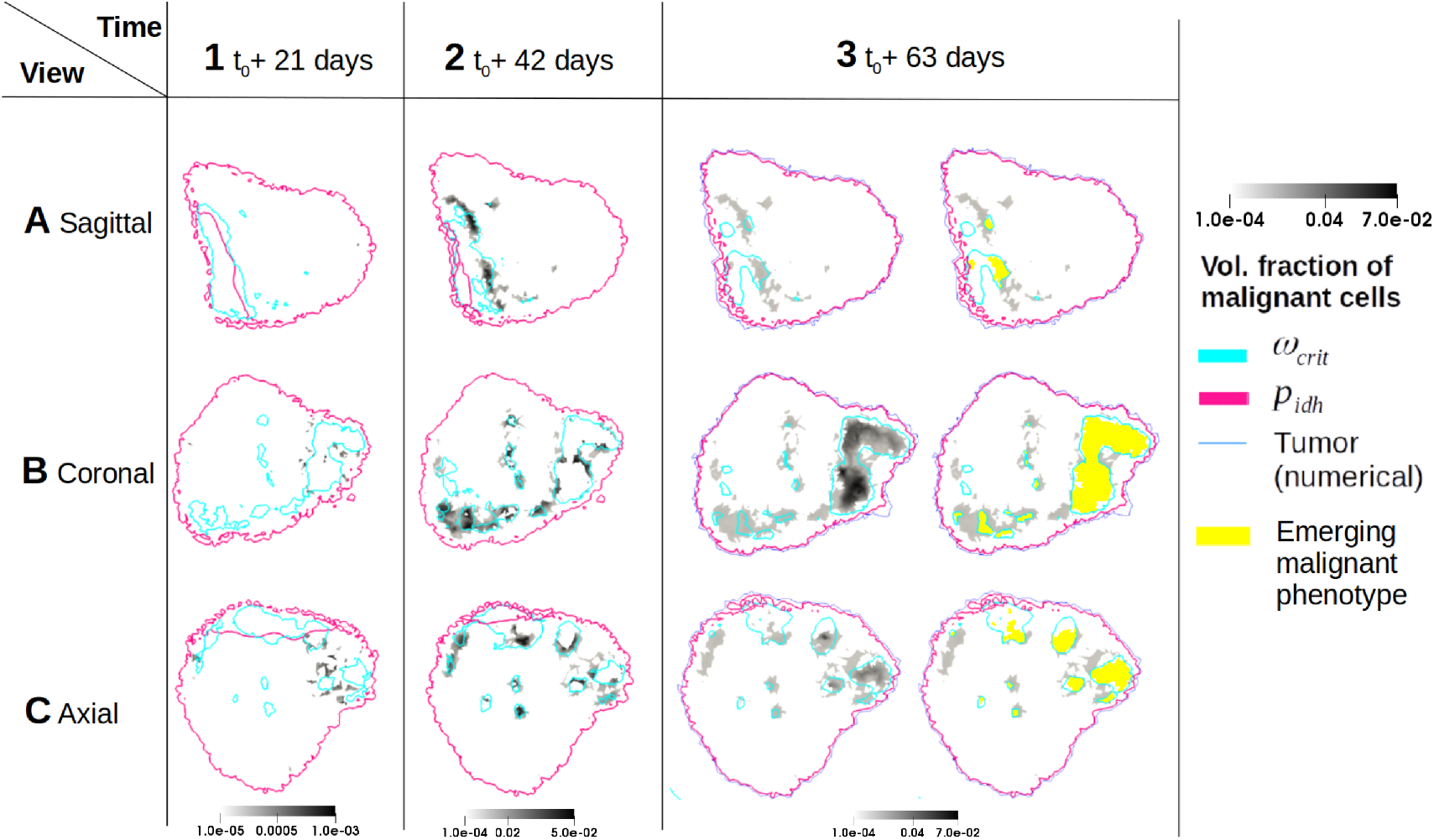
Influence of malignant phenotype in tumor growth and treatment response. **A** Sagittal view (along *x* axis). **B** Coronal view (along *y* axis). **C** Axial view (along *z* axis). **1, 2, 3** slices centered at *x* = −0.0180, *y* = −0.0104, *z* = 0.0416; volume fraction of malignant GBM cells (grey level); isoline of phenotype switch pressure *p*_idh_ (fuchsia); isoline of hypoxia threshold *ω*_crit_ (cyan); intersection of the 3 previous conditions (malignant phenotype, phenotype switch pressure and hypoxia) (yellow). **1** at *T*_0_ +21 days; **2** at *T*_0_ + 42 days after 3 cycles of RT-TMZ treatment; **3** at *T*_0_ + 63 days after 6 cycles of RT-TMZ treatment. The malignant phenotype switch is dependent of two concomitant phenomena: a high mechanical pressure (≥ *p*_idh_) inside a hypoxic environment. During the 3 first cycles of RT-TMZ treatment, Vol_mal_ is multiply by 50 fold, during the 3 last cycles Vol_mal_ is multiply by 10 fold. The details of the zones affected by phenotype switch are shown Fig.8.

##### Malignant phenotype in tumor growth and treatment response

Studies [78, 29] suggest that this phenotype switch results from an increase in internal stress, denoted by ‘tensional homeostasis’, coupled with an hypoxic environment. We proposed to model this phenomenon at the macroscale, Fig.7 shows the qualitative results of this modeling. At *T*_0_ + 21 days (Fig.7A1, B1, C1) before RT-TMZ treatment, the malignant GBM cells are rare. During the RT-TMZ treatment, the volume fraction of malignant cells Vol_mal_ initially very low ≈ 10^−5^ is 50 fold what is was during the 3 first RT-TMZ cycles, and 10 fold during the 3 latter cycles. Fig.8 shows the details of the consequences of this phenotype switch. When GBM cells are under a mechanical pressure at least equal to *p*_*idh*_ and in an hypoxic area sustained long enough (the phenotype switch is updated every 4.5 days), the development of a malignant zone occurs. Once their phenotype changed, the properties of GBM malignant cells change accordingly. The viscosity of the GBM malignant cells decreases (see Eq.27), *i*.*e*. they become more mobile. The emission of MMP degrades the ECM, so increase the intrinsic permeability of the solid scaffold (see Eq.28). Both phenomena allow for the GBM malignant fraction to rapidly invade the surrounding tissue (see Fig.8B, C).

**Figure 8:**
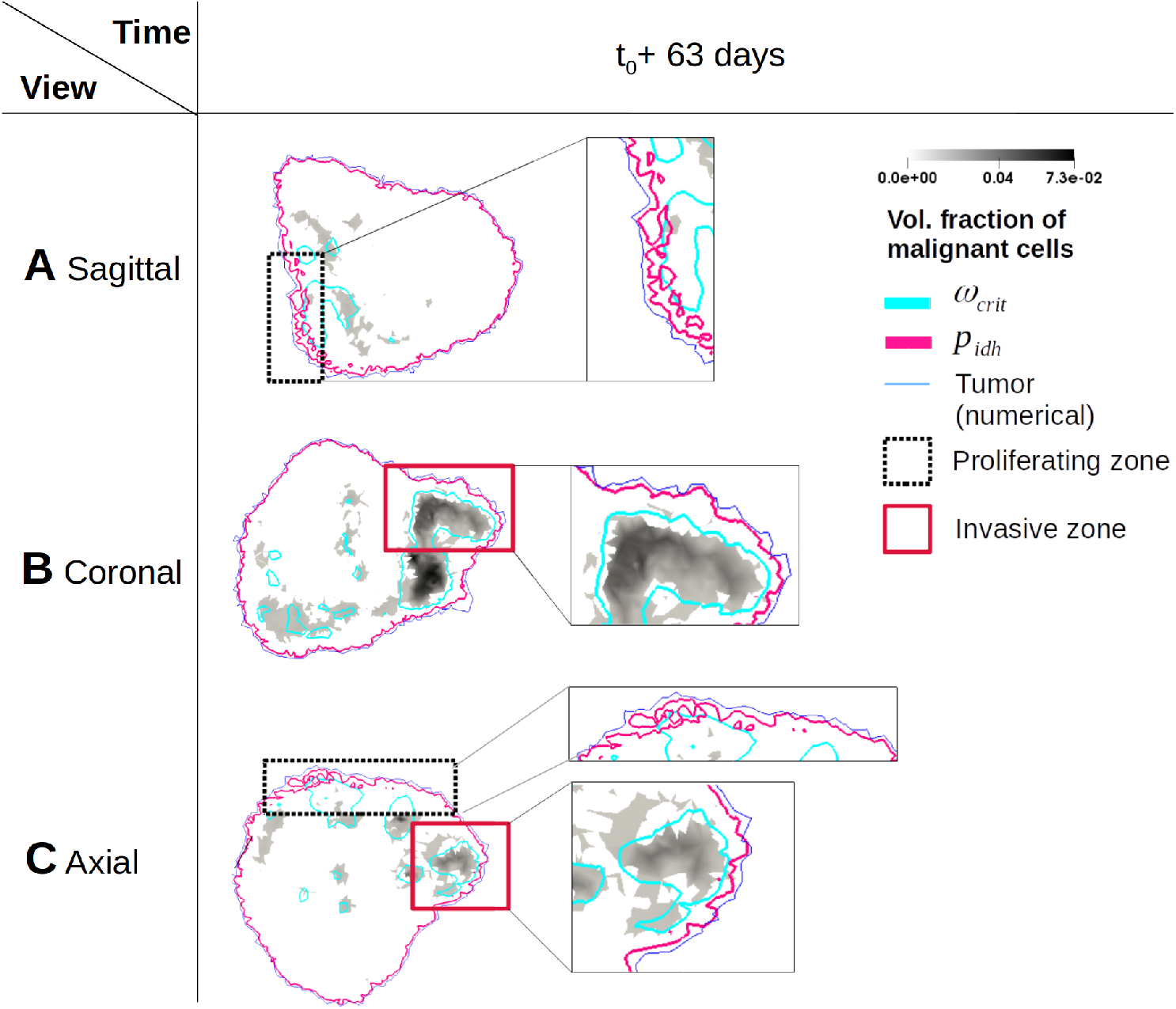
Influence of malignant phenotype in tumor growth and treatment response, focus. **A** Sagittal view (along *x* axis). **B** Coronal view (along *y* axis). **C** Axial view (along *z* axis). At *T*_0_ + 63 days after 6 cycles of RT-TMZ treatment. Volume fraction of malignant GBM cells (grey level); isoline of phenotype switch pressure *p*_idh_ (fuchsia); isoline of hypoxia threshold *ω*_crit_ (cyan); isoline *V ol*_TC_ = 0.001 (blue); proliferating zone (black dotted square); invasive zone (red square). Proliferating zones correspond to a TC growth which is not impeded by mechanical pressure, 2 zones of progression in the patient data correspond to this definition in the numerical results (one in sagittal view, one in axial view). From of point of view of the malignant evolution of the disease, numerical results show that the two mechanisms required for the phenotype switch (hypoxia and mechanical pressure) have almost no intersection in these zones. However, the zone of the sagittal view shows a beginning of an intersection. If this intersection is maintained long enough (the phenotype switch is updated every 4.5 days), it could lead to an invasive progression. Invasive zones correspond to TC migration, as the proliferation is impeded by mechanical pressure, 2 zones of progression in the patient data correspond to this definition in the numerical results (one in coronal view, one in axial view). These 2 progression zones in the patient data show a correlation between malignancy and progression in numerical results. Indeed, both mechanisms required in the phenotype switch are effective in these zones.

##### Preliminary evaluation

At *T*_0_ + 165 days, the volume of the patient tumor is 152.1 cm^3^, and the volume of the simulated tumor is 130.6 cm^3^. We obtained a simulated tumor with 14.1% of error in volume which overlaps 60.6% of the patient tumor.

Despite this important error in the preliminary evaluation, we note that the zones termed as inhibited, proliferative and invasive at the calibration time (*T*_0_ + 63 days, see Fig.6) provide relevant indications of the GBM evolution. Firstly, almost all the inhibited zones correspond to a positive response to the treatment, *i*.*e*. to a recoil of the patient tumor, or a stable zone. Secondly, all the proliferative and invasive zones correspond to a negative response to the treatment, *i*.*e*. to a progression of the patient tumor during the TMZ maintenance (see Fig.9 for details).

**Figure 9:**
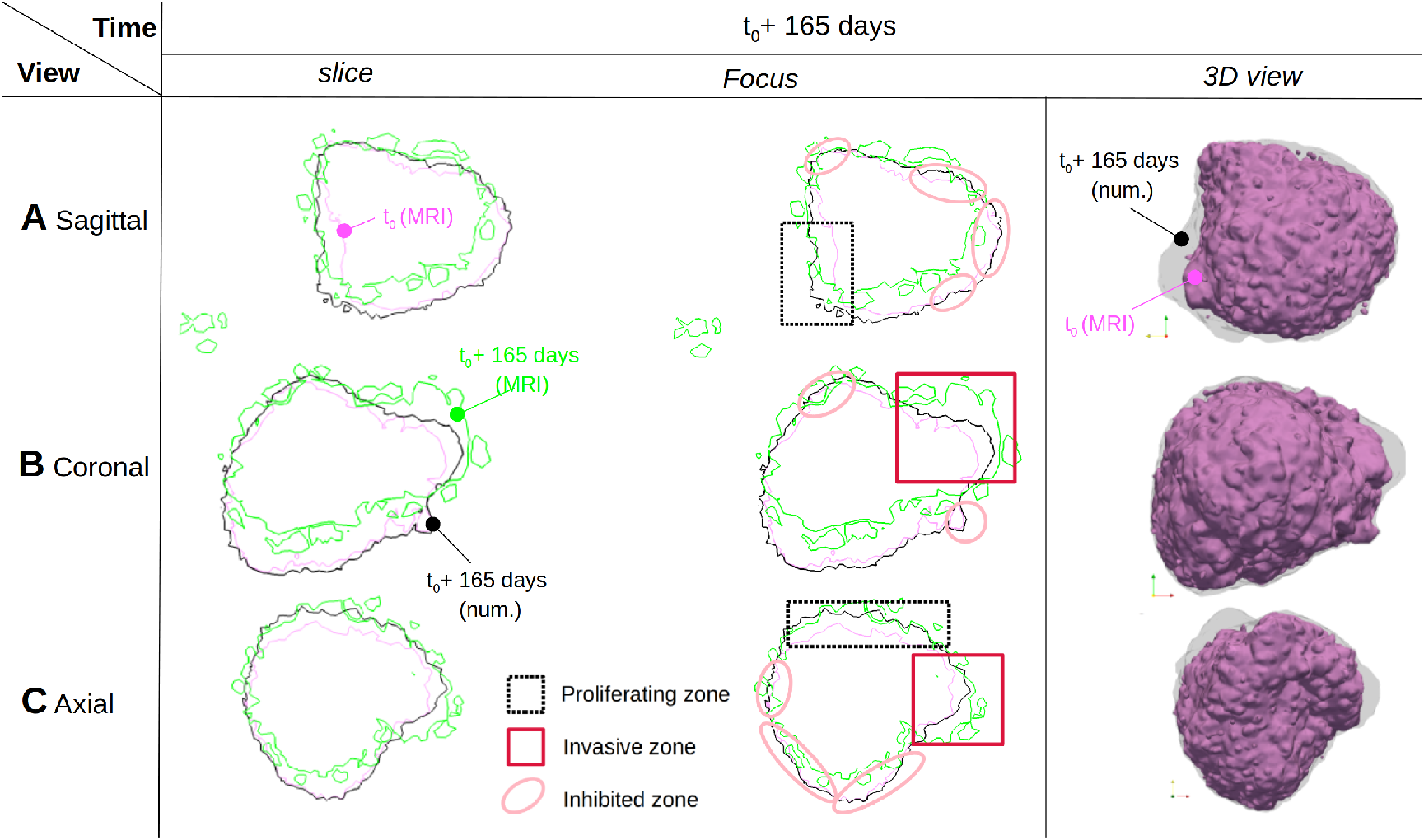
Validation against patient data at *T*_0_ + 165 days. **A** Sagittal view (along *x* axis). **B** Coronal view (along *y* axis). **C** Axial view (along *z* axis).**1** 3D isosurface Vol_TC_ = 0.001 at *T*_0_ (purple) and at *T*_0_ + 165 days (black transparent). Slices centered at *x* = −0.0180, *y* = −0.0104, *z* = 0.0416; isoline of patient data Vol_TC_ = 0.001 at *T*_0_ (purple); isoline of patient data Vol_TC_ = 0.001 at *T*_0_ + 165 days (light green) after 6 cycles of RT-TMZ treatment and 102 days of TMZ maintenance; isoline of model output Vol_TC_ = 0.001 at *T*_0_ + 165 days (black). Proliferating zone (black dotted square); invasive zone (red square); inhibited zone (pink ellipse). The numerical results that show almost no progression between *T*_0_ + 63 days and *T*_0_ + 165 days correspond to a positive response to the treatment, with a stable zone and recoil in the patient data. The numerical results that show progression correspond to a negative response to the treatment, with a large progression in the patient data. The focus column shows the comparison with the qualitative results established at the calibration outputs. Almost all the zones marked as inhibited in the simulation (tumor cells under a pressure ≥ *p*_crit_) correspond to zones with a positive response to the treatment in the patient data (no progression or recoil). The zones marked in the simulation as proliferative (neither mechanical inhibition nor hypoxia) or invasive (sustained pressure and hypoxia) correspond to a negative response to the treatment.

#### 5. Conclusions

In this study we proposed to model a patient-specific non-operable glioblastoma. The disease was first modeled within a porous medium, pre-calibrated for brain tissue in [41] by the same authors of this study. We hypothe-sized that two phenomena drive the malignant evolution of the disease: hypoxia and cell-ECM signaling. To assess patient-specific measurement, we adopted an image-informed framework. The same clinical imaging dataset (MRI methods and segmentation), at two time points, was used to initialize and calibrate the parameters. The first point was the pre-operative checkpoint and the second was performed after 6 cycles of concomitant radiochemotherapy. A last subset of parameters, which do not belong to brain tissue material properties and can not be assess by imaging, was fixed by clinical and experimental literature. After calibration, we obtained a simulated tumor with a 5.0% error in volume, comparatively to the patient tumor, and which overlaps 81.4% of the patient tumor. After 165 days, the model results were evaluated against a third dataset, we obtained a simulated tumor with a 14.1% error in volume, comparatively to the patient tumor, and which overlaps 60.6% of the patient tumor.

Qualitatively, we showed that the mechanical inhibition of the tumor growth describe well the stable zones of the patient tumor, and can partially reproduce the progression zones of the patient tumor. Thanks to the evaluation data, we showed that the mechanical inhibition corresponds to a positive response of the patient tumor to the treatment. We also showed that the proliferative and invasive zones, the latter being determined by our hypothesis of coupled high mechanical pressure and hypoxia, correspond to a negative response to the treatment.

We showed that our modeling of the GBM phenotype switch behave accordingly to the experimental findings. It has been shown that an ECM stiffer than usual brain ECM is correlated with GBM cells proliferation *and* migration [78, 79]. The same phenomena are reported under compressive stress and hypoxic environment [29]. ECM stiffness and compressive stress are linked, as in a proliferative environment, a stiffer matrix will provokes a higher internal stress. Moreover, there is no contradiction between an inhibiting pressure threshold and an internal stress, coupled with hypoxia, which provokes a malignant phenotype switch. This suggests it exists a window of mechanical signaling where GBM can dramatically evolved. Before the phenotype switch, GBM cells produce a stiffer, cross-linked, ECM. This stiffening, accompanied by the GBM proliferation, increase the internal pressure. If the pressure undergone by the GBM reaches the threshold *p*_idh_ and the level of oxygen is pathologically low (threshold *ω*_crit_), the affected GBM cells change their phenotype. They become more mobile, which is translated at the macroscale by a reduction of the dynamic viscosity of two orders. They also acquire an anaerobic metabolic pathway, which allow for escaping an hypoxic environment by metabolising lipids [80].

However, this study applies to only one patient. The response to RT-TMZ treatment and TMZ maintenance is known to be patient-specific, and without other patients or time points before the beginning and during the treatment, we cannot provide the calibration of the 3 parameters of the treatment. Therefore, we only aim to a possible model of this disease, *via* porous mechanics and mechano-biology. Apart from the application of this model to new patients, two leads are available to improve this proposition. Firstly, the parameters specific to the patient’s cell line could be pre-calibrated by exploiting the *in vitro* results of [77]. A digital twin of encapsulated tumor spheroids with a multiphase poromechanical model was validated in [25], by the same authors of the present article. In [25], the spheroids contained mouse colon carcinoma cell line, but this dual experimental setup/digital twin can be reproduced with glioma cell line. Secondly, the addition of diffusion tensor imaging method, which allow for retrieving the white matter fiber direction, would grant the access to an anisotropic permeability. This imaging method is currently a promising lead for modeling the heterogeneous progression of glioblastoma [23, 81, 82].

This study is only a first step of the inclusion of poromechanics in image-informed glioblastoma models, we hope the community will find it inspiring.

## Appendix A.

### Supporting information: solution’s sensitivity on the ROI size

Dirichlet conditions are prescribed at the ROI boundary:

- No displacement
- Fixed pressure
- Fixed oxygen level
- No necrosis

The sensitivity of these boundary conditions is evaluated on tumor evolution. After 18 days simulated, we compare the capillary pressure of the tumor phase *p*^*th*^ at each voxel of the domains with four margin sizes: 1.52 ± 0.2cm, 1.77 ± 0.3cm, 2.27 ± 0.3cm, 2.45 ± 0.4cm, denoted margin 1, 2, 3 and 4 respectively. These margins defined 4 computational domains Ω_*i*_ ∈ [1, 4] respectively. These domains contain 392 k, 425 k, 465 k and 511 k tetrahedra respectively. The larger domain Ω_4_ is used as the reference. The RMSE, without normalization, is computed as follows:

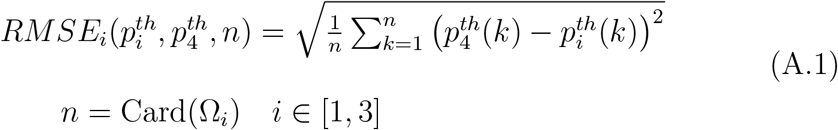

The domain evaluated is defined by the tumor cells volume fraction Vol_TC_ and its threshold is Vol_TC_ ≥ 10^−3^. This threshold corresponds, *via* l’Eq.26 for *S*^*t*^ and the range of value for the porosity, to a pressure difference of 1.4 ± 0.1 Pa. Therefore, we consider that the RMSE in Pa presented Eq.A.1 with a value above 1.4 Pa is not negligible. At day 18, the RMSE between margins 1 and 4 reaches 0.4 Pa, the RMSE between margins 2 and 4 reaches 0.17 Pa and the RMSE between margins 3 and 4 reaches 0.1 Pa. Therefore, we consider that the boundary conditions of the domain Ω_3_ have a negligible influence on the numerical solution.

**Figure A.10:**
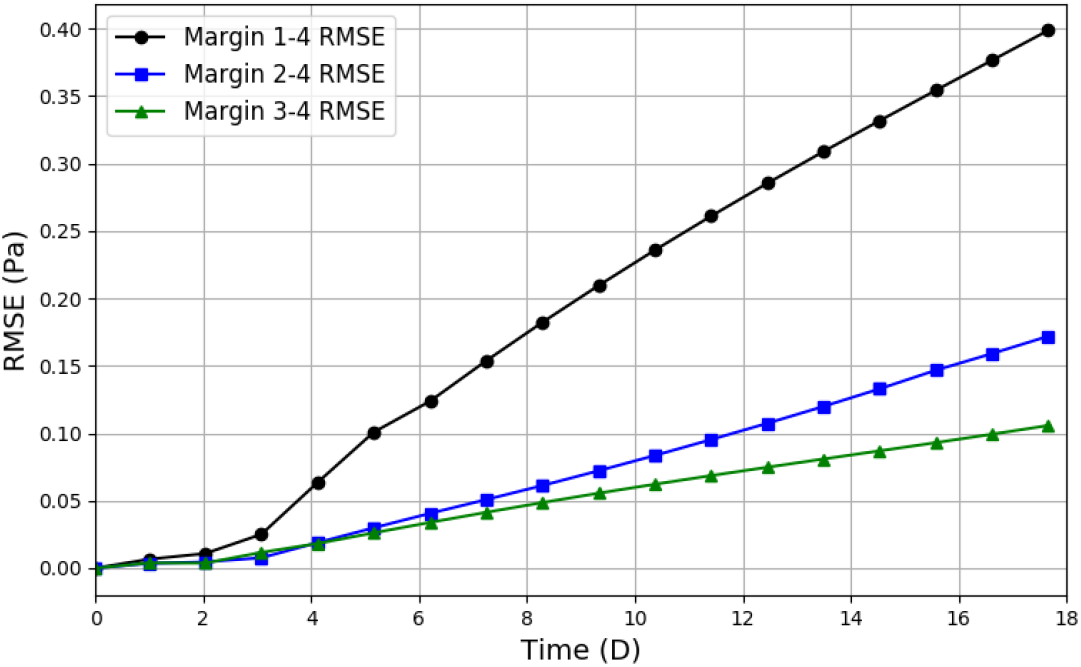
Influence of the Dirichlet boundary distance on the tumor evolution. RMSE between margins 1 and 4 (black line, circle marker); RMSE between margins 3 and 4 (green line, triangle marker). To be acceptable, the RMSE should remain below 1.4 Pa. At day 18, the RMSE between margins 1 and 4 reaches 0.4 Pa, the RMSE between margins 2 and 4 reaches 0.17 Pa, and the RMSE between margins 3 and 4 reaches 0.1 Pa

## Ethics

Informed consent was obtained from the subject involved in the study (RITC foundation for the STEMRI trial grant number RECF1929)

## Data access

The codes used in this article are available at https://github.com/SUrcun/GBM_mecano_bio. The clinical data used in this article are available upon request to the corresponding author.

## Contribution

SU: Conceptualization, Methodology, Software, Validation, Investigation, Writing – original draft, Visualization. DB: Methodology, Software, Writing - Review & Editing. PYR: Conceptualization, Investigation, Writing - Review & Editing. WS: Conceptualization, Writing - Review & Editing, Supervision. VL: Methodology, Data Curation, Validation, Investigation, Writing - Review & Editing. SPAB: Conceptualization, Writing - Review & Editing, Supervision. GS: Conceptualization, Methodology, Investigation, Writing - Review & Editing, Visualization.

## Competing interest

The authors declare that they have no known competing financial interests or personal relationships that could have appeared to influence the work reported in this paper.

## Funding

FNR-ANR

## Acknowledgement

SU thanks Tanguy Duval, Ruairidh Howells and Meryem Abbad Andaloussi for preparing the patient datasets and their segmentations. The results presented in this paper were carried out using the HPC facilities of the University of Luxembourg [83] (see https://hpc.uni.lu).

## Notes

### Competing Interest Statement

The authors have declared no competing interest.

